# PRIMITI: a computational approach for accurate prediction of miRNA-target mRNA interaction

**DOI:** 10.1101/2024.04.26.591419

**Authors:** Korawich Uthayopas, Alex G. C. de Sá, Azadeh Alavi, Douglas E. V. Pires, David B. Ascher

**Author notes:** To whom correspondence should be addressed to D.B.A.

## Abstract

Current medical research has been demonstrating the roles of miRNAs in a variety of cellular mechanisms, lending credence to the association between miRNA dysregulation and multiple diseases. Understanding the mechanisms of miRNA is critical for developing effective diagnostic and therapeutic strategies. miRNA-mRNA interactions emerge as the most important mechanism to be understood despite their experimental validation constraints. Accordingly, several computational models have been developed to predict miRNA-mRNA interactions, albeit presenting limited predictive capabilities, poor characterisation of miRNA-mRNA interactions and low usability. To address these drawbacks, we developed PRIMITI, a PRedictive model for the Identification of novel MIRNA-Target mRNA Interactions. PRIMITI is a novel machine learning model that utilises CLIP-seq and expression data to characterise functional target sites in 3’-untranslated regions (3’-UTRs) and predict miRNA-target mRNA repression activity. The model was trained using a reliable negative sample selection approach and the robust extreme gradient boosting (XGBoost) model, which was coupled with newly introduced features, including sequence and genetic variation information. PRIMITI achieved an area under the receiver operating characteristic (ROC) curve (AUC) up to 0.96 for a prediction of functional miRNA-target site binding and 0.96 for a prediction of miRNA-target mRNA repression activity on cross-validation and an independent blind test. Additionally, the model outperformed state-of-the-art methods in recovering miRNA-target repressions in an unseen microarray dataset and in a collection of validated miRNA-mRNA interactions, highlighting its utility for preliminary screening. PRIMITI is available on a reliable, scalable and user-friendly web server at https://biosig.lab.uq.edu.au/primiti.

## Introduction

Recent evidence strongly suggests that non-coding RNAs, a type of RNA that does not encode proteins, play critical roles in a variety of cellular processes, such as cell differentiation and apoptosis [1–3]. MicroRNAs, abbreviated as miRNAs, are small non-coding RNAs that regulate gene expression by partially complementary base pairing with binding sites in 3’-untranslated regions (3’-UTR) of target messenger RNA (mRNA) [4]. This process generally results in translational suppression or mRNA degradation [2–4]. Over half of all protein-coding genes are estimated to be regulated by at least one miRNA [5], highlighting the importance of miRNA functions in molecular mechanisms. Dysregulation of several miRNAs has been associated with the development of multiple diseases, such as cancers, and cardiovascular and neurological disorders [6–8].

This has led to the increasing popularity and development of personalised miRNA therapeutics, such as novel biomarkers and medications [9, 10]. Given this scenario, a range of miRNA-based drugs has been currently undergoing clinical trials [9, 10]. Despite its innovation in personalised medicine, the development of miRNA-based medical treatments is still constrained by a lack of comprehensive understanding of a miRNA targeting mechanism.

Accordingly, to gain a better understanding of miRNA activities, several experimental validation approaches for identifying transcripts targeted by miRNAs have been developed [11–16], including procedures that utilise RNA-seq or microarrays to assess changes in target expression driven by miRNA overexpression or downregulation [11, 12], or CrossLinking-ImmunoPrecipitation sequencing (CLIP-seq) to confirm the direct molecular interactions between miRNAs and mRNA target sites [13–16]. Nevertheless, large-scale experimental exploration is not feasible due to time, funding constraints and uncertainty of this process.

Consequently, several computational approaches emerged to accurately and scalably predict miRNA-target interactions, aiming to allow the creation of better diagnostic tools and treatments, and to improve miRNA-target mRNA repression characterisation. They include state-of-the-art methods such as miRanda [17, 18], RNA22 [19], PITA [20], DIANA-microT-CDS v5 [21], TargetScan [22], and miRTarget [12, 23]. A broad overview of the evolution of these methods is described in Supplementary Materials.

Despite the development effort put forward by these current methods, model usability is still greatly constrained by two main distinct factors. First, there is a limitation on sensitivity coming from CLIP-seq data [24], which is generally obscured by background noise. This usually results in an experimental failure to capture a significant number of miRNA-target interactions and reliable non-interactions. Second, there is a lack of useful characteristics defining functional miRNA-target site interactions that impair target prediction accuracy. To overcome this challenge, further improvement needs to be introduced, including not only a way to effectively select the negative samples but also incorporating characteristics that define proper functional miRNA-target interactions and non-interactions.

In this context, iLearn, a versatile feature extraction tool designed for encoding DNA, RNA, and protein sequences, offers a comprehensive set of sequence-derived physicochemical descriptive variables (or features), known as iFeatures [25, 26]. Proven to enhance model performance in various biological problems [27, 28], iLearn presents a promising approach for enhancing miRNA-target prediction accuracy. Additionally, Genome-Wide Association Studies (GWAS) have been extensively conducted globally to collect comprehensive information on human genetic variation [29]. Single nucleotide polymorphisms (SNPs), the most commonly studied genetic variation, have been observed to have an impact on miRNA repression, providing an opportunity for better characterising miRNA-binding sites [30]. The integration of iFeatures and SNPs into predictive models has the potential to provide valuable insights into the landscape of miRNA-mediated regulation.

We proposed PRIMITI, a PRedictive model for Identification of novel MIRNA-Target mRNA Interactions, to gather these insights and address the main drawbacks in current alternative methods. A machine learning model was implemented in PRIMITI to model the patterns of miRNA-mRNA interactions based on CLIP-seq and gene expression experiments [11–16]. This model incorporates more than 150 features to characterise miRNA-target site interactions, including newly introduced iFeatures and SNP features. A negative selection strategy is also employed to provide more reliable no-interactive (i.e., negative) samples.

With these attributes, PRIMITI encompasses two prediction modes, PRIMITI-TS (Target Site) and PRIMITI-TM (Target mRNA). Whereas the former was developed to predict potential miRNA-target sites that bind directly to each other, the latter was built to identify potential functional miRNA-mRNA repression activity on a particular miRNA and mRNA. In order to enhance the accessibility of our model, we provided a user-friendly web server platform for exploring human miRNA-target interactions, which is available at https://biosig.lab.uq.edu.au/primiti. We believe that with these characteristics PRIMITI would greatly contribute to a better understanding of miRNA roles, the creation of accurate biomarkers, and the development of effective treatments for a range of diseases.

## Materials and methods

### PRIMITI general workflow

The proposed methodological pipeline for both PRIMITI-TS and PRIMITI-TM comprises six primary steps, which are summarised in Figure 1. First, the datasets were collected and curated from multiple sources and experimental types [11–16]. The following step involved feature engineering for the miRNA-target site data, which resulted in the generation of 154 features. Statistical analysis was then used to examine the differences between target-site and non-target-site samples for each feature. Next, a forward stepwise greedy feature selection method [31] was utilised to choose the minimal but the most effective set of features for training the machine learning model. As a result, PRIMITI-TS was primarily developed to prioritise functional miRNA-target binding sites. Due to the fact that a single mRNA might have several target sites complementary to a particular miRNA, PRIMITI-TM was constructed using confidence scores for each target site generated by PRIMITI-TS as inputs to estimate the probability of functional miRNA suppression of mRNA targets.

**Figure 1.**
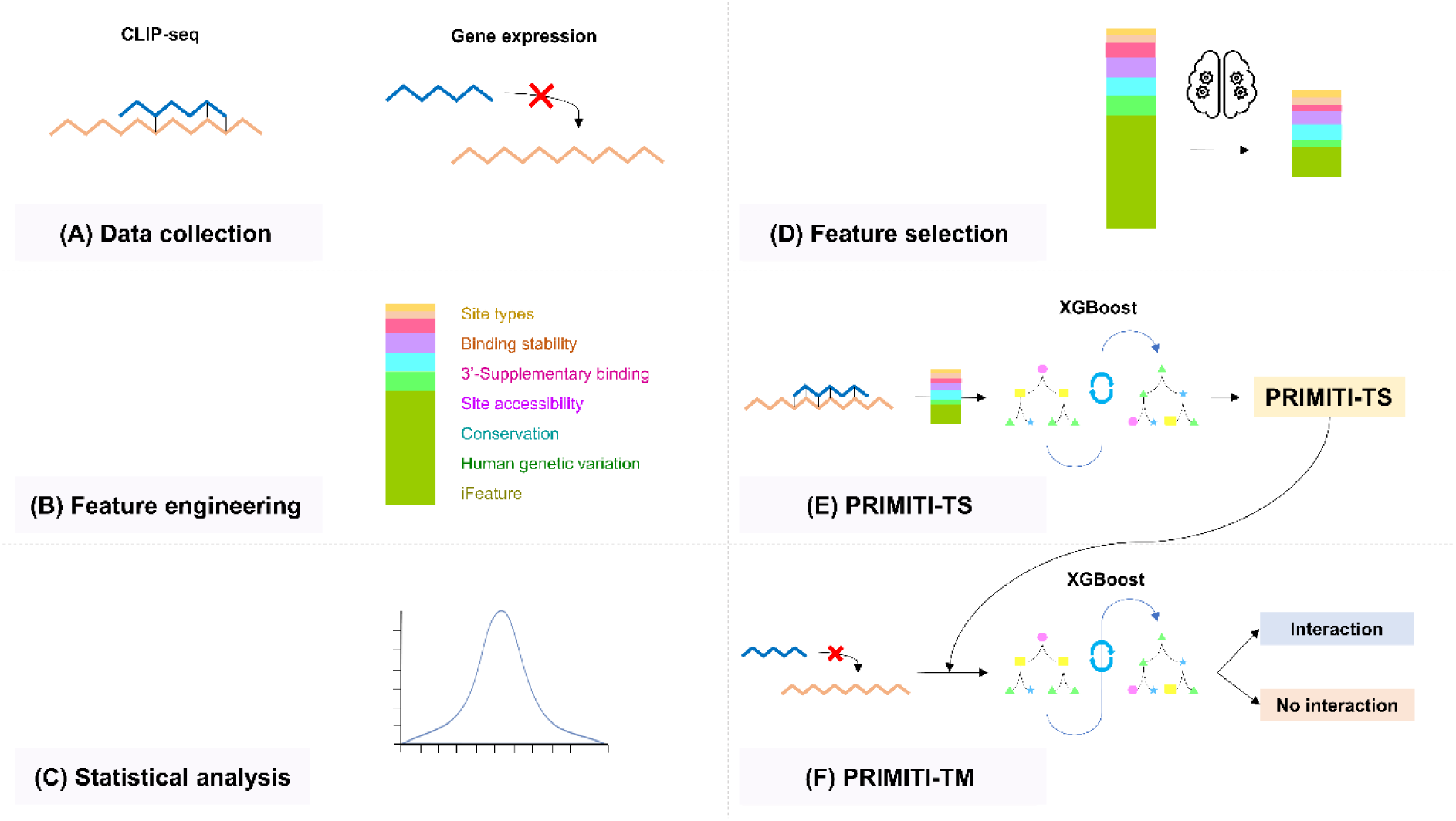
PRIMITI workflow: miRNA-target mRNA interaction prediction. The development of PRIMITI is divided into six steps: (A) Data collection – experimental CLIP-seq and gene expression data are gathered from multiple sources, (B) Feature engineering – seven types of features are generated to characterise miRNA-target site interactions, (C) Statistical analysis – Mann-Whitney U test and Chi-square test were used to analyse the binding between miRNAs and target sites, (D) Feature selection – relevant features were determined using forward stepwise greedy feature selection, (E) PRIMITI-TS model training and evaluation – the extreme gradient boosting (XGBoost) algorithm was employed to predict miRNA-target site interactions, (F) PRIMITI-TM model training and evaluation – prediction results from PRIMITI-TS were used as features to construct a final XGBoost model for prioritising potential miRNA-mediated mRNA repression.

### Data Collection and Preprocessing

As aforementioned, PRIMITI is divided into two prediction modes, PRIMITI-TS and PRIMITI-TM. Consequently, they differ during data acquisition, which is described next. The list of experimental data is provided in Table S1.

In PRIMITI-TS, data collection starts by curating miRNA-target site interactions (i.e., positive samples). In this step, CLIP-seq data was acquired from different sources [13–16] to be used in PRIMITI. The verified miRNA-canonical target site interactions constitute 6,190 positive samples between 300 miRNAs and 3,331 mRNA transcripts from 3,147 genes. In terms of non-interactions (i.e., negative samples), 504,159 negative samples were extracted from unverified canonical miRNA-target site interactions from the same CLIP-seq data. Nevertheless, because of the limited sensitivity of CLIP-seq, interpreting unconfirmed interactions as non-interactions may result in the production of poor-quality negative data. Although this practice is commonly used in alternative methods [12, 22], we aimed to improve this aspect on negative data. To increase the quality of negative samples, negative sample selection was therefore introduced. 451,413 reliable negative samples were selected by comparing them to a collection of experimentally validated miRNA-target mRNA interactions from miRTarbase [32] and Tarbase [33]. Any miRNA-target mRNA pair identified in this list was considered to have an interaction with each other and was subsequently removed from the negative sample dataset. Given the high-imbalance ratio between interactions and non-interactions, a random under-sampling procedure [34] was used to create a dataset with 1:1 balanced data with miRNA-mRNA target interactions and non-interactions. Finally, high-throughput sequencing of RNA isolated by crosslinking immunoprecipitation (HITS-CLIP) data [13] was utilised as an additional external test set. 98 miRNA-target site samples and 7,240 miRNA-non-target site samples were derived after using the same validation procedure of CLIP-seq data.

In PRIMITI-TM, on the other hand, miRNA-target mRNA repressions were extracted from miRTarget’s RNA-seq data [12]. By mapping repressed genes with more than 40% reduction in gene expression as target mRNAs, a total of 2,351 miRNA-target mRNA interactions were found between 25 miRNAs and 1,511 genes. In contrast, a set of negative samples was created from genes with unaffected expression levels, ranging from 100% to 110% when compared to a negative control sample without miRNA transfection. With these thresholds, we obtained 6,749 miRNA-non-target mRNA pairs involving 25 miRNAs and 3,966 genes. To validate PRIMITI-TM and compare it to alternative methods, microarray data from Linsley *et al*. [11] was utilised. After normalisation and definition of functional miRNA-mRNA repression activities, 596 miRNA-mRNA interactions and 10,233 miRNA-mRNA non-interactions were observed between 7 miRNAs and 1,279 genes.

### Feature generation

We carried out an extensive literature review, searching for features that could assist in the characterisation of miRNA-target site interactions. As a result, 154 features are developed and generated to delineate a miRNA-target site interaction for PRIMITI-TS. The set of features consists of 4 canonical site types, 4 binding stability, 13 site accessibility, 7 3’-supplementary bindings, 16 conservation, 18 human genetic variations, and 92 iFeature descriptors. They are briefly introduced as follows. A complete description of the feature generation is provided in Supplementary Material and Methods, and in Table S2.

● ***Canonical site types***. Canonical site type features are related to different suppression efficacy in terms of mRNA fold change and can be divided into four classes: 6-mer, 7-mer-m8, 7-mer-A1, and 8-mer [4].
● ***Binding stability.*** The stability of the miRNA-induced silencing complex (miRISC) complex formed by miRNA, mRNA, and Argonaute2 (Ago-2) protein, is referred to as the binding stability. This has been substantially correlated with miRNA targeting efficacy [35] and, therefore, used in PRIMITI.
● ***Site accessibility.*** Accessibility to a binding site is considered a critical factor while determining the targeting specificity and translational repression activity [36].
● ***3’-supplementary binding.*** During the binding process, and after the seed region has initiated a conformational change of miRISC, the last half of the 3’-region of miRNA is exposed for additional interactions with the target mRNA [37, 38]. This supplementary interaction is regarded as a 3’-supplementary pairing and is mostly occurring at the 3’-supplementary regions [39].
● ***Conservation.*** Conservation, in turn, is a useful indicator for prioritising functional target sites [5, 39], as genetic regions coding for functional miRNA binding sites have been proven to evolve at a slower rate due to evolutionary constraints to the conserved functional regions [5].
● ***Human genetic variation.*** Additionally, human genetic variants were implemented and used to assist in the characterisation of miRNA-binding sites. A presence of single nucleotide (SNPs) and disease-related SNPs on each location in a target site is considered.
● ***iFeatures.*** iLearn is a newly presented Python toolkit that implements a comprehensive set of descriptive variables (or features), named as iFeatures, to encode structural and physicochemical information of nucleic acid or peptide sequences [25, 26]. It has been widely used in multiple fields such as a prediction of mRNA subcellular localisation or protein-protein interaction [27, 28]. It is important to highlight and stress that this is the first time iFeatures are used to characterise miRNA-target site interactions.

Statistical analysis for this set of features has been discussed in detail in Supplementary Material and Methods and thereafter provided in Supplementary Results.

### PRIMITI-TS Model training

A range of supervised machine learning (ML) algorithms was independently assessed [40], including random forest, extreme gradient boosting (XGBoost), extremely randomised trees, adaptive boosting, gradient boosting, artificial neural networks, decision trees, k-nearest neighbours, support vector machines and gaussian processes.

Among them, XGBoost [41] outperformed the nine other algorithms when training PRIMITI-TS with the complete feature set (see Table S3). We believe the reason for this is because of XGBoost’s main characteristics. XGBoost is a highly optimised decision tree-based ensemble machine learning algorithm that has advantages in efficiency, flexibility, and parallelisation. Additionally, it can manage missing data internally, obviating the need to alter the data explicitly prior. This technique combines the results of numerous weak tree-based classifiers to obtain increasingly precise overall predictions. Consequently, we have decided to keep XGBoost for the remaining parts of the PRIMITI-TS pipeline.

Internal cross-validation approaches, as well as external validation approaches with independent blind test sets, were performed to evaluate the performance of PRIMITI-TS using different metrics, including the Area Under the Receiver Operating Characteristic Curve (AUC), Balanced accuracy (bACC), F1-score (F1), Matthew’s Correlation Coefficient (MCC), Precision, Recall, and Specificity. An independent blind test set comprised samples separated from the training dataset prior to model training. Detailed information about the performance metrics employed is available in Supplementary Materials and Methods.

### PRIMITI-TS Feature selection

Given the current set of 154 features, a forward stepwise greedy feature selection procedure is performed in PRIMITI-TS aiming to select the best combination of features while maintaining the model’s simplicity [31, 42, 43]. Greedy feature selection starts with a set of zero features and then selects the feature that contributed the most to the predictive performance at that iteration. The predictive performance of the XGBoost model [41] in terms of MCC across a 10-fold cross-validation procedure (Figure S1) was used to quantify the quality of each feature [40]. Iteratively, the following features are chosen similarly and added one by one in the current set. In the end, a set of 22 features was selected based on the analysis trading-off between complexity and predictive performance on 10-fold cross-validation across greedy feature selection iterations (Figure S1 and Table S4). This set contains two site types, one binding stability, one 3’-supplementary binding, three site accessibility, four conservation, two human genetic variations, and nine iFeature features [25]. Overall, this selection shows that all types of features are important in predicting the interactions between miRNAs and target mRNAs. Hence, the final PRIMITI-TS model was trained using these 22 features with the XGBoost algorithm.

### PRIMITI-TM Model training

PRIMITI-TS was previously developed to predict a functional binding interaction between miRNAs and target sites. However, in certain real-world scenarios, researchers are more concerned with miRNA-mediated regulation of mRNA, which is a biological consequence of physical interactions between miRNA and multiple target sites in one mRNA. Previous work used this information to improve the understanding of the association of miRNAs and prostate cancer [44–46] and to develop miRNA therapeutics and biomarkers [44–46].

It is worth noting that predicting suppression is not equivalent to predicting binding sites, since a single target mRNA may host several binding sites and not all physical binding sites can result in miRNA-mediated gene silencing. When evaluating repressive interactions, the likelihood of each putative site must be considered collectively. PRIMITI-TM has been proposed to prioritise miRNA-target mRNA interactions. PRIMITI-TS will first calculate the likelihood of all potential canonical binding sites for each miRNA-mRNA interaction. Then, four features based on calculated probabilities will be used to train PRIMITI-TM’s model, including: (i) the summation of all probabilities in all target locations, (ii) the probability in the highest probability site, (iii) the probability in the second highest probability site, and (iv) the probability in the third highest probability site (Figure S2). Afterwards, these features will be used to characterise experimentally verified miRNA-mRNA suppression. 5-, 10– and 20-fold cross-validation procedures and an independent blind test set were utilised to evaluate the performance of PRIMITI-TM’s model.

Overall, PRIMITI advances in the field of miRNA-target interaction by considering new aspects for characterising RNAs (e.g., iFeature and human genetic variation), the development of a negative sample selection method for choosing the reliable miRNA-target non-interactions, and a novel proposal to model miRNA-target site binding and suppression.

## Results

### Performance of PRIMITI-TS

We employed a number of cross-validation procedures to evaluate the model used in PRIMITI-TS in prioritising miRNA-target site interactions. On 5-fold cross-validation, PRIMITI-TS achieved AUC bACC, F1 and MCC values of 0.958, 0.894, 0.894 and 0.788, respectively. In addition, PRIMITI-TS yielded comparable results on 10– and 20-fold cross-validation procedures, indicating its ability to accurately predict miRNA-target site interactions (Table 1 and Figure 2A).

**Figure 2.**
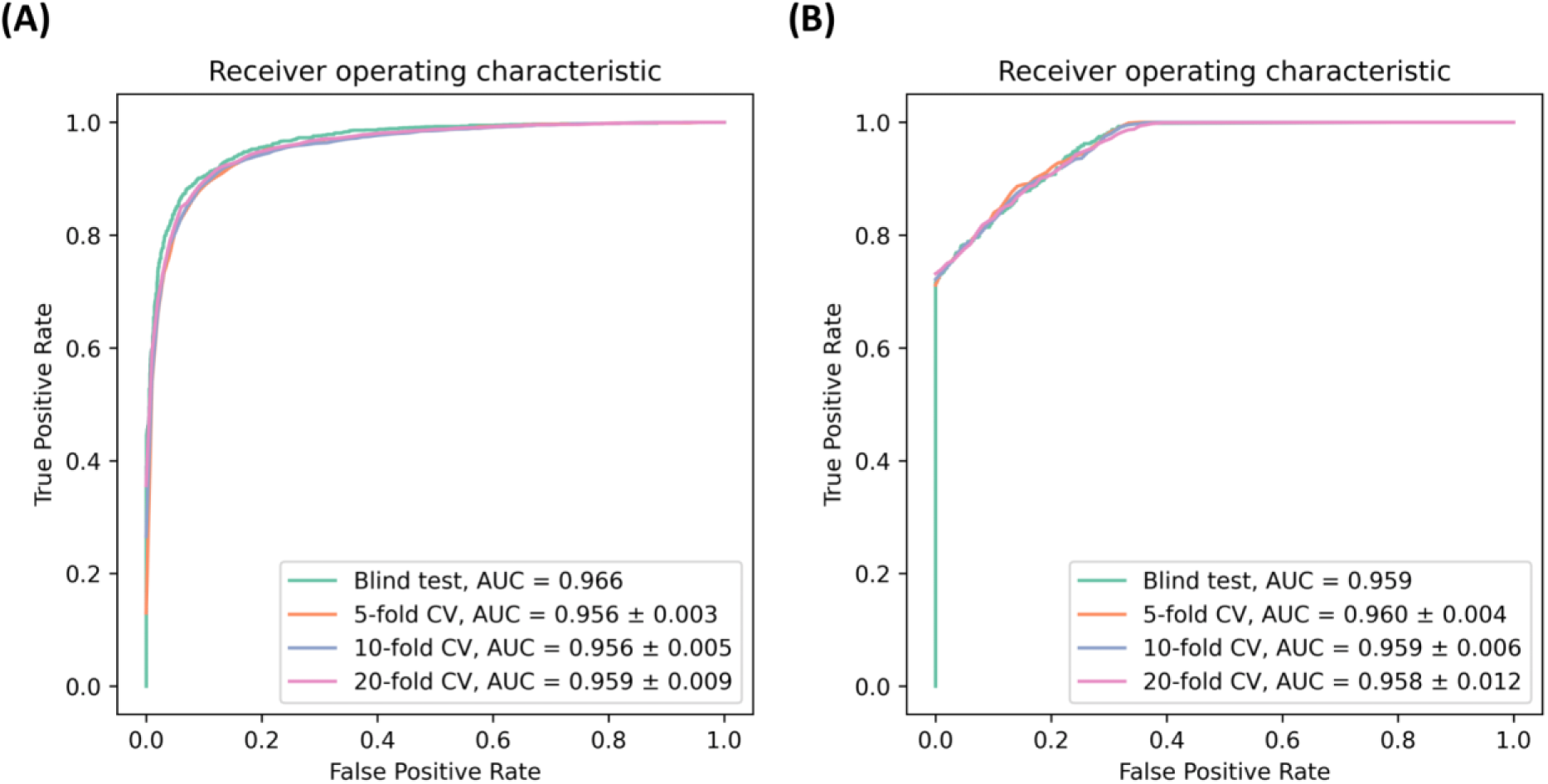
The Receiver Operating Characteristic (ROC) Curve Analysis of PRIMITI. (A) PRIMITI-TS successfully predicts miRNA-target site interactions in a blind test, 5-fold, 10-fold, and 20-fold cross-validation (AUC of 0.966, 0.956, 0.956, and 0.959, respectively). (B) PRIMITI-TM, a model trained with miRNA-mRNA repression activities, effectively prioritises potential repression activity, with AUC values of 0.959, 0.960, 0.959, and 0.958 in a blind test, 5-fold, 10-fold, and 20-fold cross-validation, respectively.

**Table 1.** The performance evaluation of PRIMITI-TS on 5-fold, 10-fold, and 20-fold cross-validation, and on a blind test set.

| Method | AUC | bACC | F1 | MCC | Precision | Recall | Specificity |
| --- | --- | --- | --- | --- | --- | --- | --- |
| 5-fold cross-validation | $0.956 \pm 0.003$ | $0.898 \pm 0.006$ | $0.897 \pm 0.006$ | $0.796 \pm 0.012$ | $0.900 \pm 0.006$ | $0.895 \pm 0.008$ | $0.900 \pm 0.006$ |
| 10-fold cross-validation | $0.956 \pm 0.005$ | $0.899 \pm 0.007$ | $0.899 \pm 0.006$ | $0.798 \pm 0.013$ | $0.901 \pm 0.013$ | $0.896 \pm 0.010$ | $0.901 \pm 0.015$ |
| 20-fold cross-validation | $0.959 \pm 0.009$ | $0.903 \pm 0.017$ | $0.903 \pm 0.018$ | $0.807 \pm 0.035$ | $0.906 \pm 0.018$ | $0.901 \pm 0.027$ | $0.906 \pm 0.020$ |
| Blind test | 0.966 | 0.904 | 0.903 | 0.809 | 0.915 | 0.892 | 0.917 |

Additionally, we conducted an assessment across an independent blind test to estimate PRIMITI-TS’ ability to generalise across unseen data. The model has achieved AUC, bACC, F1 and MCC scores of 0.965, 0.898, 0.897 and 0.797, respectively. These results are consistent with the predictive results from the employed cross-validation procedures. (Table 1 and Figure 2A).

The predictive performance of PRIMITI-TS to previously unknown data was further evaluated using HITS-CLIP data [13] (Table S5 and Figure S3) with the assessment of several classification thresholds. After this, PRIMITI reached bACC, F1, MCC, Precision, and Specificity up to 0.918, 0.148, 0.258, 0.080 and 0.846 at a cut-off of 0.900, while achieving a Recall of 0.990.

Supplementary analyses are conducted to assess the model’s generalisability and the results are presented in Supplementary Results (Table S6-S7). An analysis aims to uncover the model’s capability to generalise across different experimental data sources (Table S1) by constructing models using two datasets and evaluating their performance on a different dataset. The results indicate that all models demonstrate a notable degree of generalisation across the datasets, with an AUC of approximately 0.9. Despite a minor decline in performance, the model can accurately predict outcomes in diverse *C. elegans* cells [15] when trained on human cells [14, 16], achieving AUC, bACC, F1 and MCC scores of 0.895, 0.813, 0.803 and 0.628, respectively (Table S6).

Further analysis was performed to evaluate the model’s capability to handle newly unseen miRNAs and transcripts (Table S7). Three independent tests are being conducted using 30% of miRNAs and/or mRNAs that have been separated from the training set. Although the models’ performance is slightly affected by a lack of characterisation on new miRNAs and/or transcripts, they are still capable of effectively applying their knowledge to new data, achieving AUC, bACC, F1 and MCC values more than 0.920, 0.840, 0.830 and 0.690, respectively.

The results provided in this section and in the Supplementary Materials corroborate the robustness and accuracy of PRIMITI-TS for properly predicting unseen interactions between miRNAs and mRNA target sites.

### PRIMITI-TS’s interpretation

Model interpretability is a critical aspect for demonstrating the viability of use in a real-world situation by providing insight into how the model works [47, 48]. Thus, we have employed SHapley Additive exPlanation (SHAP) [48] to analyse the relationships between the features and model’s output, as illustrated in Figure 3. The features in this figure are listed in descending order according to the degree of impact on model prediction. We observed that the most crucial feature is the overall interaction energy estimated from IntaRNA. The miRNA-target site combination that requires low energy tends to be a functional target site, which biologically corresponds to the fact that a miRNA-mRNA duplex structure that requires lower energy is more energetically stable. Additionally, this interpretation results demonstrate that site type is a significant predictor of a functional target site. Being an 8-mer site increases the likelihood of being a functional target site, while 6-mer increases the likelihood of being a non-functional target site. This can be explained by the fact that an 8-mer site contains seven seed complementary pairings and adenine at the first position, resulting in the highest binding affinity compared to a 6-mer site with the lowest affinity. Because binding affinity correlates with repression efficiency [35], the status of the site type plays a crucial role in determining functional repression. This is consistent with a prior analysis that exhibits a significant statistical difference in the distribution of site types between positive and negative samples (Table S8).

**Figure 3.**
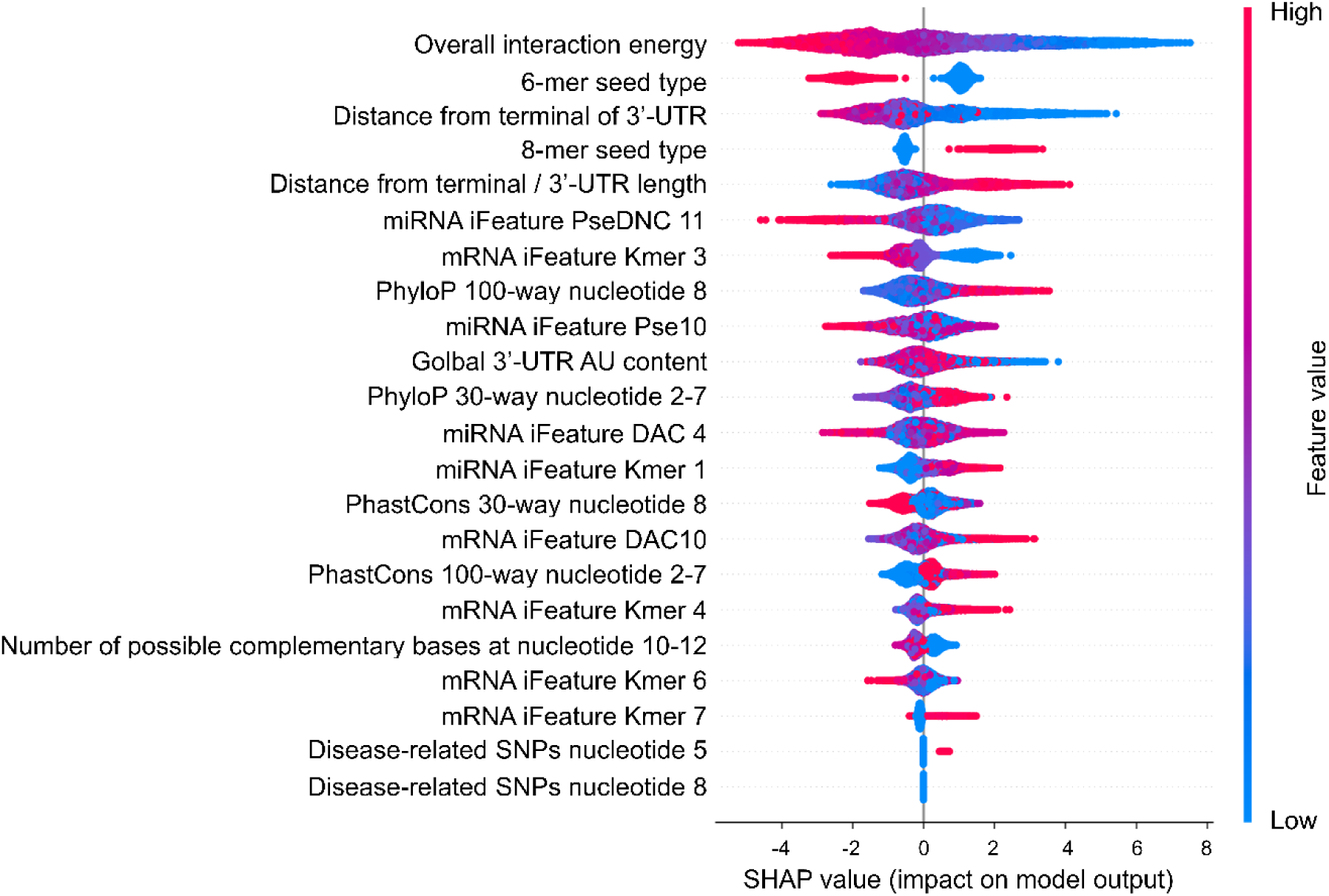
The most essential feature is an overall interaction energy, followed by 6-mer seed type and distance from the 3’-UTR terminal. The SHAP value was computed for each feature in PRIMITI-TS. The features are sorted according to their impact on the model’s prediction accuracy. Each dot denotes an interaction between a miRNA and its target site, while colours denote the values of features, with red denoting high values and blue denoting low values.

We also discovered that targets located near the 3’-UTR are more likely to be functional, which is consistent with a previous study demonstrating that putative target sites are more evolutionarily conserved in the area towards the start and end of the 3’-UTR owing to a greater degree of site accessibility [36]. Furthermore, SHAP results also indicated that single-nucleotide polymorphism (SNP) characteristics provide only a small contribution to model prediction, corroborating the prior analysis’s finding (Figure 3). We hypothesise that the limited data availability of reported disease-related SNPs limited the contribution of SNP features toward the performance [49].

### Performance of PRIMITI-TM

Similar to the evaluation process of PRIMITI-TS, a blind test, 5-fold, 10-fold, and 20-fold cross-validation procedures were used to assess the performance of PRIMITI-TM in predicting miRNA-target mRNA repression activity. On 5-fold cross-validation, PRIMITI-TM obtained an AUC, bACC, F1, and MCC values of 0.961, 0.867, 0.834, and 0.765, respectively (Table 2). Comparable performances were achieved using 10-fold and 20-fold cross-validation, respectively (Table 2). This demonstrates that PRIMITI-TM is capable of accurately predicting the suppression of miRNA-target mRNA. We conducted a blind test to determine PRIMITI-TM’s generalisation capabilities. Equivalent performance in cross-validation was achieved, with AUC, bACC, F1, and MCC values of 0.959, 0.868, 0.835 and 0.764, respectively. Receiver operating characteristics curves (ROC) of all validations are shown in Figure 2. The comparable performances in a blind test, as well as in 5-fold, 10-fold, and 20-fold cross-validation, support the robustness of PRIMITI-TM in predicting miRNA-target mRNA interactions.

**Table 2.** PRIMITI-TM performance evaluation on 5-fold, 10-fold, and 20-fold cross-validation, and a blind test.

| Methods | AUC | bACC | F1 | MCC | Precision | Recall | Specificity |
| --- | --- | --- | --- | --- | --- | --- | --- |
| 5-fold cross-validation | 0.960 ± 0.004 | 0.867 ± 0.012 | 0.834 ± 0.016 | 0.765 ± 0.021 | 0.902 ± 0.015 | 0.777 ± 0.021 | 0.957 ± 0.007 |
| 10-fold cross-validation | 0.959 ± 0.006 | 0.864 ± 0.013 | 0.830 ± 0.018 | 0.758 ± 0.026 | 0.897 ± 0.026 | 0.773 ± 0.025 | 0.955 ± 0.014 |
| 20-fold cross-validation | 0.958 ± 0.012 | 0.864 ± 0.020 | 0.831 ± 0.026 | 0.760 ± 0.037 | 0.901 ± 0.032 | 0.771 ± 0.033 | 0.957 ± 0.015 |
| Blind test | 0.959 | 0.868 | 0.835 | 0.764 | 0.896 | 0.782 | 0.954 |

To further evaluate the generalisation of PRIMITI-TM, experimental miRNA-mRNA repression activities from Linsley *et al*. [11] were employed. This dataset is notably different from the dataset used to train PRIMITI-TM in terms of an experimental approach and cell line. Numerous computational models have been presented to date for predicting miRNA-mRNA interactions. In this study, we compare the performance of PRIMITI-TM with four state-of-the-art predictors: miRTarget [12, 23], TargetScan [22], DIANA-microT-CDS v5 [21], and RNA22 [19]. While the chosen techniques are trained on a variety of datasets, none of them is trained on the Linsley *et al.* microarray [11], which would allow a fair comparison. The prediction results from respective websites were retrieved and compared. Due to the highly imbalanced dataset, we validated the findings using MCC, F1, bACC, Precision, Recall, and Specificity. The result demonstrates that selected predictors may be classified into two categories: models that prioritise precision (miRTarget, DIANA-microT-CDS v5, and RNA22) and models that prioritise recall (PRIMITI-TM and TargetScan). Among all methods, miRTarget delivers the best overall performance (MCC, F1, and bACC) (Table 3). However, miRTarget’s model was built at the cost of a poor recall rate. This implies that a substantial proportion of functional miRNA-target mRNA interactions will be likely to be predicted as non-interactions. In contrast, our model captures the majority of positive samples, providing the greatest recall of any model. This contributes to the applicability of PRIMITI-TM for initial interaction screening. Additionally, PRIMITI outperforms TargetScan, the other recall-focused model, on all metrics, demonstrating a higher predictive potential of PRIMITI-TM (Table 3). We recommend combining PRIMITI-TM with several precision-focused predictors to efficiently identify miRNA-target interactions with high confidence.

**Table 3.** Validation of PRIMITI-TM based on Linsley microarray data.

| Model | MCC | F1 | bACC | Precision | Recall | Specificity |
| --- | --- | --- | --- | --- | --- | --- |
| **PRIMITI** | 0.022 | 0.108 | 0.523 | 0.059 | **0.681** | 0.364 |
| **TargetScan** | 0.006 | 0.103 | 0.507 | 0.056 | 0.661 | 0.352 |
| **miRTarget** | **0.111** | **0.164** | **0.602** | **0.102** | 0.418 | 0.786 |
| **DIANA-microT-CDS v5** | 0.052 | 0.121 | 0.540 | 0.084 | 0.220 | **0.860** |
| **RNA22** | -0.041 | 0.078 | 0.455 | 0.044 | 0.336 | 0.575 |

Additionally, our findings were corroborated using a dataset of 759,764 experimentally validated miRNA-target mRNA interactions obtained from miRTarbase [32] and Tarbase [33]. We select 20 well-studied miRNAs with the highest number of validated targets. For each miRNAs, we randomly select 100 positive samples and 100 negative samples, resulting in a balanced dataset of 4,000 miRNA-target interactions. The process was repeated for 10 replicates. The performance of PRIMITI-TM was compared with that of miRTarget [12, 23], TargetScan [22], DIANA-microT-CDS v5 [21], and RNA22 [19] (Table 4). The results suggest that both miRTarget and PRIMITI-TM are capable of achieving the greatest overall performance, as measured by MCC, F1, and bACC. However, when recall is taken into account, PRIMITI-TM attained a significantly higher score (0.466) than miRTarget (0.176) (Table 4).

**Table 4.** A validation based on a dataset of experimentally validated miRNA-target mRNA interactions retrieved from miRTarbase and Tarbase databases.

| Model | MCC | F1 | bACC | Precision | Recall | Specificity |
| --- | --- | --- | --- | --- | --- | --- |
| **PRIMITI** | $0.108 \pm 0.012$ | $0.510 \pm 0.010$ | $0.553 \pm 0.006$ | $0.564 \pm 0.007$ | $0.466 \pm 0.013$ | $0.641 \pm 0.008$ |
| **TargetScan** | $0.079 \pm 0.015$ | $0.471 \pm 0.010$ | $0.538 \pm 0.008$ | $0.551 \pm 0.010$ | $0.411 \pm 0.012$ | $0.666 \pm 0.012$ |
| **miRTarget** | $0.158 \pm 0.015$ | $0.283 \pm 0.009$ | $0.552 \pm 0.005$ | $0.709 \pm 0.021$ | $0.176 \pm 0.006$ | $0.928 \pm 0.006$ |
| **DIANA-**<br>**microT-CDS**<br>**v5** | $0.049 \pm 0.011$ | $0.034 \pm 0.004$ | $0.505 \pm 0.001$ | $0.725 \pm 0.058$ | $0.017 \pm 0.002$ | $0.993 \pm 0.002$ |
| **RNA22** | $0.005 \pm 0.008$ | $0.103 \pm 0.003$ | $0.501 \pm 0.002$ | $0.511 \pm 0.016$ | $0.058 \pm 0.002$ | $0.945 \pm 0.003$ |

In summary, the main reason for the great predictive performance achieved by PRIMITI is related to its reliable data (including the miRNA-target site non-interactions, i.e., the negative samples), the advances in characterising both miRNA-target site binding and repression, and the novelties while modelling target-site interactions with machine learning, including a new way to summarise the target site information as features to predict miRNA-target repression. As a consequence, our findings – which are based on the implemented methodology – indicate that PRIMITI is an excellent tool for pre-screening miRNA-target interactions.

## Online web interface

PRIMITI is accessible to the public in the form of a web server with a user-friendly interface via https://biosig.lab.uq.edu.au/primiti/. PRIMITI’s Prediction and Results pages are demonstrated in Figure 4. In Figure 4A, users must supply a list of miRNAs or a single miRNA following the universal miRBase format. Users must also provide a list of mRNAs or a single mRNA within their transcript IDs, where an Ensembl Transcript (ENST) format should be preferably used. Details on alternative input formats are listed at https://biosig.lab.uq.edu.au/primiti/help. PRIMITI will then process the inputs, providing a result table (Figure 4B). Each pair of miRNA-target interactions is accompanied by a confidence score for the miRNA-target mRNA repression determined by PRIMITI-TM, indicating the likelihood of functional mRNA-miRNA repression. Details of each miRNA-target binding site are additionally provided (Figure 4B).

**Figure 4.**
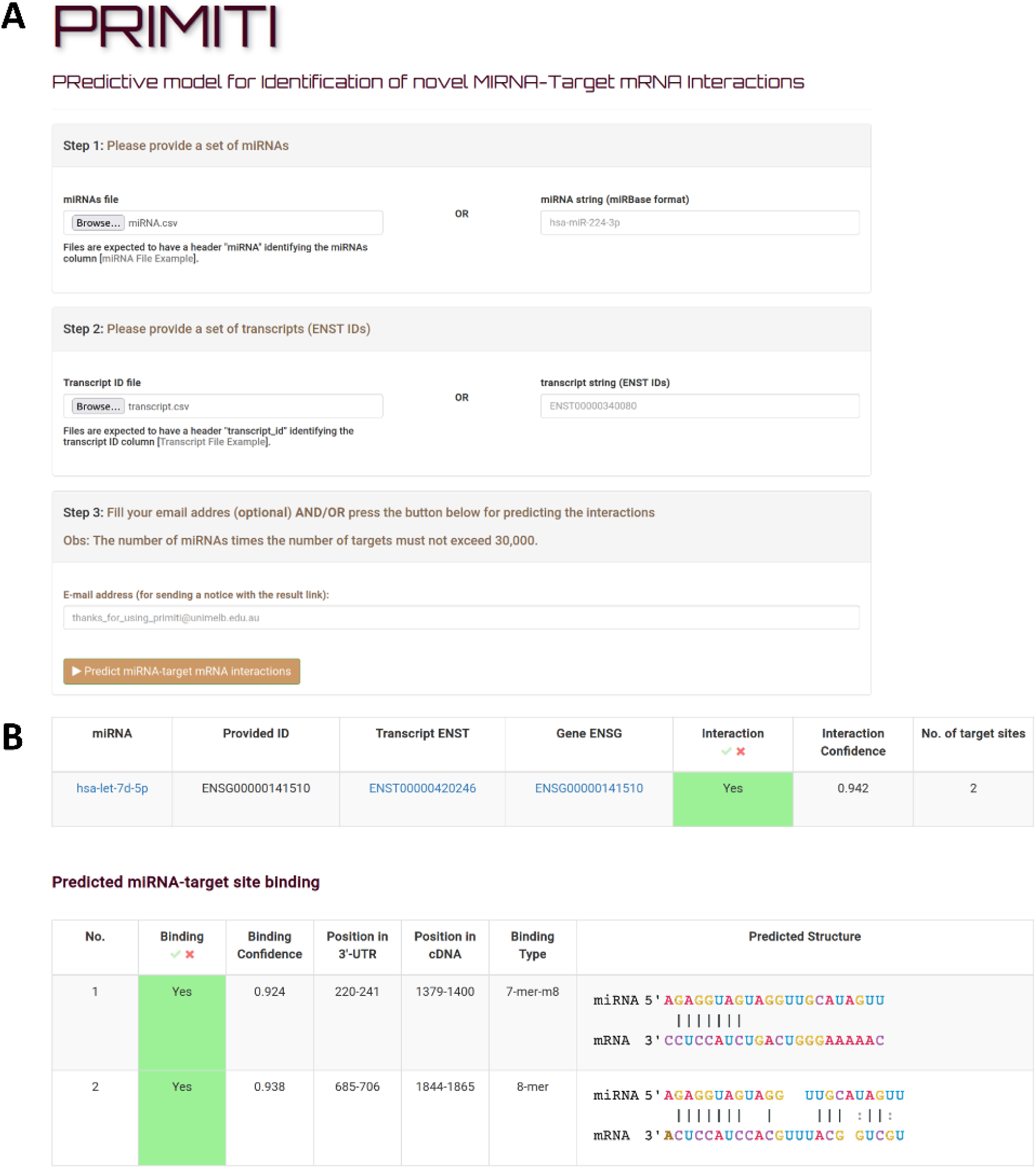
PRIMITI web server interface. (A) The PRIMITI web server requires an input file containing a list of miRNAs in miRBase format and transcripts or gene identifiers. In the case of gene identifiers, the longest transcripts will be selected for a prediction. (B) The result of PRIMITI is provided in tables. Prediction scores for PRIMITI-TS and TM will be given to each binding site and transcript, in which a higher score indicates a higher probability of interaction.

In addition, all the experimental data used to train, (cross-)validate and test PRIMITI’s model can be downloaded via its web server at https://biosig.lab.uq.edu.au/primiti/data. Furthermore, all code and data related to PRIMITI can be downloaded or accessed through GitHub at https://github.com/winkoramon/PRIMITI-miRNA-target-mRNA-interactions.

## Discussion

MicroRNAs play a crucial role in controlling gene expression networks, which are responsible for a wide range of cellular processes [1–3]. Distinct miRNA profiles are present in every cell type, precisely influencing cellular phenotype. The dysregulation of miRNA functions is associated with various human diseases [6–8]. Exploring the functions of miRNA provides valuable knowledge about disease mechanisms, which helps in the discovery of biomarkers and the development of innovative therapies [9, 10]. The introduction of high-throughput sequencing techniques has improved our understanding of the activities of miRNAs, with a particular focus on their effects on target mRNAs [12–16]. With this progress, computational approaches have arisen as a solution to the limitations of traditional methods, enabling efficient analysis of high-throughput transcriptomic data. These approaches allow for large-scale investigation of miRNA-mediated interactions [12, 17–23].

In this work, we proposed and developed PRIMITI, a novel machine learning model for predicting miRNA-target mRNA repression. PRIMITI identifies patterns between miRNA-mediated repression of mRNA in 3’-UTR from CLIP-seq and gene expression (RNA-seq or microarray) data [11–16]. In PRIMITI’s method, we have also introduced negative sample selection to filter out low-quality negative samples and improve the reliability of the training dataset (Table S9). Moreover, genetic variation and sequence-derived information are implemeted as a way to improve the characterisation of miRNA-target site binding in PRIMITI-TS. Sequence-derived information is carried out by iFeatures, which are a set of features generated by iLearn’s sophisticated scheme for encoding nucleic sequences into numerical features [25, 26]. Human genetic variation was brough by human-based SNPs and is another area that may host useful information for the miRNA repression characterisation [30, 49]. As a result, PRIMITI succeeds in predicting miRNA-mediated repression with an AUC and MCC up to 0.96 and 0.80 for a prediction of functional miRNA-target site binding and up to 0.96 and 0.76 for a prediction of miRNA-target mRNA repression activity. This predictive performance is achieved by PRIMITI on 10-fold cross-validation over the training set and independent blind test. The model also achieves a good performance in predicting repression in an unseen microarray dataset and collection of validated miRNA-mRNA interactions, yielding the largest number of validated miRNA-target repressions accurately predicted as repressed, when compared to state-of-the-art methods, demonstrating the utility for preliminary screening (Table 3-4). PRIMITI is made available either in a GitHub repository (https://github.com/winkoramon/PRIMITI-miRNA-target-mRNA-interactions) or a web server https://biosig.lab.uq.edu.au/primiti). PRIMITI offers a number of benefits for the large-scale identification of miRNA-target interaction by providing valuable experimental guidance for the validation of predicted targets. By prioritising computationally selected candidates, researchers can optimise their experimental efforts, leading to more targeted and successful validations, as well as improving the pipeline’s efficiency and cost-effectiveness. PRIMITI’s model also provides a comprehensive understanding of miRNA-target site binding, including binding probability and binding structure (Figure 4B).

### Limitations

There are a few limitations associated with this work. Firstly, a lack of training data may limit the generalisation of our model. PRIMITI was trained on a relatively small dataset, considering that there are over 2000 unique human miRNAs [50], 100,000 unique transcripts, and a distinct set of cell types in the human body. Despite this limitation, our model still demonstrates reasonable predictive capabilities when trained on human data [14, 16] and tested on drastically different *C. elegans* data [15] (Table S6). However, we anticipate that incorporating additional data will considerably improve the model’s predictive performance and, as a consequence, generalisation across numerous cell lines and species.

In addition, the sensitivity of the experiment may have an impact on the model accuracy, particularly in regard to negative samples. Obtaining reliable negative samples is crucial for the development of a robust machine learning model. However, due to limited sensitivity, it becomes challenging to acquire true negative samples, as some positives can be misidentified as negatives. To address this challenging aspect, we implemented a negative sample selection approach to increase the reliability of negative samples, which resulted in a significant performance improvement (Table S9). Nonetheless, an enhanced experimental method with increased sensitivity has the potential to further improve the overall performance of our proposed model.

### Feature generation

In this work, PRIMITI-TS have implemented new sets of features, iFeature and single nucleotide polymorphisms (SNPs). iFeature, a nucleic acid sequence encoding scheme provided by iLearn, has demonstrated significant enhancements in various machine learning models across diverse research domains [27, 28]. Our analysis of the model’s performance indicated a substantial improvement when integrating iFeature into PRIMITI-TS (Table S10). The 5-fold cross-validation results showed an increase in the Matthews correlation coefficient (MCC) from 0.71 to 0.80. This finding also suggests that employing advanced encoding schemes can effectively aid in the characterisation of target sites. In subsequent studies, it would be beneficial to investigate the potential advantages of employing other encoding schemes, such as RNABERT [51], to further improve the performance of the model.

Furthermore, within the framework of PRIMITI-TS, we hypothesised that leveraging human genetic variations could aid in the characterisation of miRNA-binding sites. It has been observed that SNPs occur across verified miRNA-targeted mRNA sequences, with some of them contributing to dysregulated gene expression regulation and the progression of disease [30]. Based on this supporting evidence, we integrated a set of SNPs and disease-related SNPs from miRNASNP v3 [49] into the PRIMITI-TS model. However, our analysis did not reveal any significant differences in performance between models trained with and without SNP features (Table S10). This unexpected outcome implies that additional research is needed to determine the significance of SNP traits in miRNA-target mRNA interactions and to evaluate the extent of human SNP data coverage in miRNA targets. For example, we intend in future research to evaluate the benefits of exploring alternative representations for SNPs. We also plan to incorporate multiple data sources to increase the quantity of variation data.

## Conclusion

miRNAs play an essential role in post-transcriptional gene regulation through complementary base pairing with target sites in mRNAs. Therefore, identifying target sites in mRNAs and target mRNAs is an essential task for leading not only to an improvement of diagnostic tools and treatments but also to a better understanding of the whole human epigenetic regulation. In this study, PRIMITI was developed to identify miRNA-target mRNA interactions through machine learning modelling, incorporating novelties in terms of the types of employed features (i.e., iLearn-based and SNP-related features) to better characterise functional miRNA binding sites, as well as negative sample selection to provide a more reliable training set for a machine learning model. PRIMITI was assessed internally and externally validated, resulting in a robust predictive performance in terms of area under the ROC curve (AUC), Matthew’s correlation coefficient, Recall (or sensitivity) and specificity. PRIMITI’s predictive results indicate a great generalisation, providing the ability to correctly identify miRNA-target mRNA repression. Different from other alternative methods, PRIMITI is made available as a robust and user-friendly predictive web server at https://biosig.lab.uq.edu.au/primiti/. In future work, we plan to improve PRIMITI by ensembling it with other alternative methods and models. We trust this would extend PRIMITI’s current predictive performance, especially on the precision metric.

## Code and data availability

The data used to train and test PRIMITI-TS and PRIMITI-TM and other supplementary data is available at https://biosig.lab.uq.edu.au/primiti/data. The source code for PRIMITI is available at https://github.com/winkoramon/PRIMITI-miRNA-target-mRNA-interactions.

## Highlights

● miRNAs play an essential role in post-transcriptional gene regulation through complementary base pair binding with mRNA target sites.
● PRIMITI yields a new machine learning model (ML) model to identify miRNA-target mRNA interactions.
● PRIMITI incorporates novelties in the characterisation of functional miRNA binding sites and also in providing more reliable training sets for its respective ML model.
● PRIMITI achieved great predictive performances under cross-validation, blind test, and independent test sets, indicating its robustness in identifying miRNA-target mRNA repression.
● PRIMITI was made available as a user-friendly web server at https://biosig.lab.uq.edu.au/primiti/.

## Acknowledgements

We would like to thank Carlos Rodrigues for his assistance on web server deployment.

