## Supplementary document 1 for "PRIMITI: a computational approach for accurate prediction of miRNA-target mRNA interaction"

### SUPPLEMENTARY LITERATURE REVIEW

#### Evolution of computational methods to predict miRNA-target interactions

miRanda [1, 2] is one of the earliest methods that recognizes target sites using basic position-weighted local alignment and free energy estimated from ViennaRNA RNAfold [3] while filtering targets with low conservation scores. In this way, miRanda is able to predict target mRNAs at the cost of being unable to foresee non-conserved areas. RNA22 [4], in turn, attempts to solve the miRNA-target interaction problem in a completely different way. Its solution involves identifying the distinct characteristics of functional target sites before pairing with candidate miRNAs. Nevertheless, this method suffers from a low predictive performance for identifying mRNA-miRNA interactions.

Later, PITA [5], which stands for Probability of Interaction by Target Accessibility, eliminated the need for a cross-species sequence conservation filter, enabling the discovery of non-conserved targets. The model employed by PITA introduced a site accessibility characteristic, which is represented by the energy required to unpair intramolecular pairing in mRNAs. This approach makes the target sites accessible for miRNA binding. However, a strong dependence on thermodynamic stability calculated by ViennaRNA RNAfold [3] often results in inaccurate structural predictions.

Aiming to go beyond the predicted performance of the aforementioned methods, DIANA-microT-CDS v5 [6], TargetScan [7], and miRTarget [8, 9] have incorporated additional characteristics related to a degree of conservation [10] and 3'-supplementary pairing [11] of functional target sites to increase the model predictive performance on identifying potential interactions between target mRNAs and miRNAs. These innovative characteristics considerably improve model practicality. As a result, these models have been applied in a wide range of scientific investigations [12-14]. For instance, a recent study implemented TargetScan, miRTarget, and miRanda to identify target genes in The Cancer Genome Atlas (TCGA) dataset, uncovering a number of tumour-suppressor miRNAs and the promise for miRNA replacement treatment [12].

### SUPPLEMENTARY MATERIAL AND METHODS

#### Data collection - PRIMITI-TS

***Positive sample curation.*** A complete list of datasets analysed in our work is provided in Table S1. High-throughput CrossLinking-ImmunoPrecipitation (CLIP) sequencing approaches have been applied to identify regions in the mRNA transcripts that bind with the Argonaute (Ago) protein [15-18]. The Ago protein is loaded with a single-stranded miRNA, forming microRNA-Induced Silencing Complexes (miRISC), which in turn bind target mRNAs via seed complementarity. CLIP-seq data was acquired from Helwak *et al.* [17], Grosswendt *et al.* [16], and Kozar *et al.* [18]. The verified miRNA-canonical target site interactions constitute 6190 positive samples between 300 miRNAs and 3331 mRNA transcripts from 3147 genes.

***Reliable negative sample generation.*** The canonical miRNA-target sites are reported to be associated with significant repression in several experiments due to the presence of a seed complementary sequence [7]. They play a critical role in the initial pairing between miRNA and mRNA seed region (nt 2 - 7), and a stable formation of miRNA-mRNA duplex. Hence, the canonical target site is generally the primary focus of target prediction models. For each experiment, we compiled a list of potential binding interactions between miRNAs and canonical target sites identified in the transcripts that reported binding activity. Pairs of miRNAs and target sites that were not present in positive samples were regarded as non-interactions and were thus categorised as negative samples. Relying on this biological characteristic, 504,159 negative samples were produced from unverified canonical miRNA-target site interactions. However, given CLIP-seq's limited sensitivity, the strategy of generating negative samples by considering unconfirmed pairings to be non-interacting may create low-quality negative samples. In this work, we implemented a negative sample selection strategy to enhance the reliability of the negative samples. All negative samples were validated with a list of 759,764 experimentally validated miRNA-target mRNA interactions from miRTarbase [19] and Tarbase [20]. A pair of miRNA and target mRNA detected on the list were assumed unreliable negative samples since it had evidence of repression activity, and were thus excluded from the negative sample dataset. As a consequence, a reliable set of 451,413 negative samples was obtained. A general under-sampling procedure [21], was used to create a dataset with a 1:1 positive to negative ratio, meaning a balanced ratio between miRNA-mRNA target interactions and non-interactions, respectively. This method was employed on confirmed miRNA-target site interactions yielding a final positive:negative balanced dataset of 12,322 miRNA-target site pairs, in which 30% of experimentally confirmed miRNA-target sites from were isolated from the training set and retained for evaluation purposes.

***Independent test sets.*** Apart from a variation of CLIP-seq that contains a ligation step in PRIMITI-TS training dataset, HITS-CLIP data, conducted by Boudreau *et al*. [15], was utilised as an independent test set. The potential miRNA-target interactions were extracted, however, without a ligation step in HITS-CLIP, exact and unambiguous relationships between miRNAs and target sites cannot be determined. A markedly different technique to extract potential miRNA-target site interactions was adopted based on the original article [15]. Experimentally established RISC-associated target sites that contain a seed complementary sequence, an indicator of canonical binding, were employed to search for matching seed sequences in all members of the top five most prevalent miRNA families. Only members of the top five most prevalent families were chosen to provide more plausible miRNA-target site interactions and, therefore, minimise false positives due to the increased likelihood of highly expressed miRNA targets being reported in HITS-CLIP. As a result, 177 possible miRNA-target interactions were identified. To strengthen the robustness of interactions, we incorporated experimentally established miRNA-mRNA interactions from miRTarbase [19] and Tarbase [20] to verify a set of potential interactions. When interactions were found in both databases, they were considered to be reliable miRNA-target site interactions, and were thus included in a positive sample dataset. Simultaneously, miRNA-non-target site interactions, known as negative samples, were constructed from pairs of miRNA and non-overlapping target sites in the collection of transcripts found in CLIP-seq. This approach guarantees that the transcripts used for generating negative samples are expressed in the cell. We validated the absence of interactions using the same set of experimentally confirmed interactions in miRTarbase [19] and Tarbase [20]. Given this validation procedure, a dataset of 7,338 miRNA-target site pairings was established. It comprises 98 miRNA-target site samples and 7,240 miRNA-non-target site samples.

#### Data collection - PRIMITI-TM

On the other hand, in order to train the PRIMITI-TM, a model for prioritising functional miRNA-mRNA repression activity, a dataset containing expression information was necessary. miRNA-target mRNA repressions were extracted from RNA-seq data reported in miRTarget [8, 9]. The translational repression activities involving 25 miRNAs were profiled in terms of an expression change. The experiment started with a transfection of miRNA into HeLa cell lines, followed by RNA-seq. The experiment was duplicated to control the experimental variations. The same approach as in the original article [8, 9] was used to extract functional miRNA-target mRNA repression signatures. Significantly repressed genes with more than 40% reduction in gene expression will be regarded as target mRNAs. Accordingly, a total of 2,351 miRNA-target mRNA interactions were discovered between 25 miRNAs and 1,511 genes. In contrast, a set of negative samples was created from genes with unaffected expression levels, ranging from 100% to 110% when compared to a negative control sample without miRNA transfection. Finally, we obtained 6,749 miRNA-non-target mRNA pairs involving 25 miRNAs and 3,966 genes.

Raw microarray data from Linsley *et al*. [22] was retrieved from the NCBI GEO database (accession GSM156522, GSM156523, GSM156524, GSM156532, GSM156547, GSM156548, GSM156580) to validate PRIMITI-TM and compare it to other state-of-the-art models. The raw microarray data was normalised in Bioconductor using the Robust Multi-Array average (RMA) approach [23]. In these experiments, a single miRNA was transfected into HCT116 cell lines for overexpression. Genes with statistically significant differential expression were considered to receive functional miRNA-mRNA repression activities. 596 miRNA-mRNA interactions and 10,233 miRNA-mRNA non-interactions were observed between 7 miRNAs and 1,279 genes.

#### Characterising miRNA-target site interactions

To characterise miRNA-target site interactions, the sequences of miRNAs were obtained from miRBase (release 22) [24], and the transcript reference sequences were retrieved from Ensembl (release 104) via the Ensembl API [25]. If a gene name, NCBI accession number, or Ensembl Gene ID is provided instead of an Ensembl Transcript ID, Mygene API [26] will be used to map it to an Ensembl Transcript ID. If more than one transcript is related to the given accession, an Ensembl Transcript ID with the longest UTR will be chosen as it is more likely to host the largest number of target sites. Pre-calculated PhyloP and Phastcons conservation scores for 100-way and 30-way multiple alignments can be directly retrieved from UCSC Genome Browser using genomic coordinates [27, 28]. Single Nucleotide Polymorphisms (SNP) and disease-related SNPs in 3'-UTR were extracted from the miRNASNP-v3 database in order to describe the miRNA-target site interactions in terms of human genetic variations [29]. 759,764 confirmed miRNA-mRNA interactions were retrieved from miRTarbase [19] and Tarbase [20]. Data generated by public miRNA-target predictors, miRTarget [8, 9], TargetScan [7], DIANA-microT-CDS v5 [6], and RNA22 [4], was obtained from the respective online repositories from these tools. For direct comparison, the target transcript IDs were mapped to Ensembl transcript IDs.

#### Feature generation

***Canonical site types***. Canonical site types can be divided into four classes: 6-mer, 7-mer-m8, 7-mer-A1, and 8-mer [30]. The 6-mer target site contains nucleotides that are perfectly complementary to mature miRNA position 2-7, whereas the 7-mer-m8 site contains nucleotides perfectly matched with position 2 to 8. 7-mer-A1 is similar to 6-mer as both are complementary to miRNA position 2-7, but differs in the presence of an adenine (A) nucleobase at the first position of mRNA. 8-mer presents an exact match to positions 2 to 8, with the 'A' nucleotide at the first position. Due to the pocket in a Argonaute-2 (Ago-2) protein that exclusively binds to adenine, the presence of the adenine is related with increased stability of the miRISC complex, resulting in an improved suppression activity [31]. A study in *Drosophila* [32] revealed that each site type results in different suppression efficacy in terms of mRNA fold change, with 8-mer being the most effective site, followed by 7-mer-m8, 7-mer-A1, and 6-mer.

***Binding stability.*** The stability of the miRISC complex formed by miRNA, mRNA, and Ago-2 protein, referred to as the binding stability, is substantially correlated with miRNA targeting efficacy [33]. Greater stability would cause increased occupancy at a target site and thus increased repression of a target mRNA [33]. The amount of energy required for miRNA-target site interactions, generated from RNA-RNA interaction prediction models, is generally used to represent the stability. The less energy required to form the complex, the more stable the binding is. miRNA-target interaction prediction models employ a variety of strategies to predict the most stable RNA-RNA duplex structure, resulting in different calculated binding stability. Recent research [34] has benchmarked 14 state-of-art RNA-RNA duplex structure prediction methods. Among all methods compared by Lai and Meyer [34], IntaRNA achieved the highest predictive performance. The method has one primary advantage over alternative methods, in which it takes site accessibility into consideration when predicting the most energetically stable structure. Hence, in this work, we employed IntaRNA to predict the most energetically stable miRNA-target site duplex and binding stability features, including overall interaction energy, hybridization energy, and overall interaction energy of seed as features to be used to predict miRNA-target site interaction.

***Site accessibility.*** Accessibility to a binding site is a critical factor in determining the targeting specificity and translational repression activity. Grimson *et al.* [35] conducted a conservation and site-depletion analysis to ascertain the factors influencing a target-site accessibility. The results in their analysis established the effect of accessibility on miRNA repression activity and unveiled a link between an AU content of neighbouring regions (flanks) and target-site accessibility. Flanks are represented by nucleotide residues, which are located at 30 nucleotides upstream and downstream of the target site. According to them, the flanks of functional binding sites are highly enriched in A and U nucleobases, resulting in a weaker secondary structure and increased accessibility of mRNA. Not only the AU content of flanks has effects on target site accessibility, but also the target site's position in the 3'-UTR. In 3'-UTR, effective sites are preferentially located near the stop codon and poly A tail. This is because sites near the ends are more accessible due to less occlusive mRNA intramolecular interaction and are closer to the translational machinery.

***3'-supplementary binding.*** During the binding process, after the seed region has initiated a conformational change of miRISC, the last half of the 3'-region of miRNA is being exposed for additional interactions with the target mRNA [11, 36]. This type of supplementary interaction is regarded as 3'-supplementary pairing, which mostly occurs at 3'-supplementary region (nt 13-16) [37]. It significantly stabilises the miRISC and affects the target recognition [11]. Recent study revealed a large number of non-overlapped target sites for miRNAs within the same family with an identical seed, indicating a bias in target specificity caused by 3'-supplementary bindings [38]. Several characteristics, such as numbers of possible complementary bases in each region and GC content, involving 3'-supplementary pairing are included in the PRIMITI model. The richness of possible complementary bases and GC content in 3'-supplementary regions has shown to relate with higher target affinity and can bear a longer loop structure [11, 39].

***Conservation.*** Conservation, in turn, is a useful indicator for prioritising functional target sites [10, 40]. Genetic regions coding for functional miRNA binding sites have proven to evolve at a slower rate due to evolutionary constraints to the conserved functional regions [10, 40]. Thus, PRIMITI utilises two basewise conservation metrics, PhyloP and PhastCons [41, 42], calculated from 100 and 30 Vertebrate species multiple alignments, to represent the degree of conservation in a single nucleotide resolution. PhyloP and PhastCons employ a different approach to calculate the conservation level. PhastCons [42] incorporates flanking nucleotides into the calculation, increasing the metric's sensitivity for finding more conserved areas. PhyloP, on the other hand, ignores the impact of their neighbours entirely, making it more suitable for studying selection signals at a particular nucleotide [41]. The conservation scores were implemented in PRIMITI for four locations - 1$^{st}$ nucleotide, 2$^{nd}$ - 7$^{th}$ nucleotide region, 8$^{th}$ nucleotide, and 13$^{th}$ - 16$^{th}$ nucleotide region - which contribute differentially towards a binding stability [36]. For regions, median values were selected to represent the conservation level, as it is less sensitive to outliers.

***Human genetic variation.*** Additionally, human genetic variants can also be utilised to assist in the characterization of miRNA-binding sites. Genome-Wide Association Studies (GWAS) have been widely conducted worldwide to collect data on genetic variations and their associations with various traits in different individuals [43]. Single-Nucleotide Polymorphisms (SNPs), a primary focus of GWAS, have been identified across various verified mRNA target sequences [43]. Several of them result in dysregulated gene expression regulation, which contributes to disease development [44]. For instance, an rs2057482 (T>C) SNP near the miR-199a binding region had been associated with pancreatic ductal adenocarcinoma and several clinical characteristics such as tumour's size and survival rate [45, 46]. PRIMITI creates 14 features to represent SNPs and disease-related SNPs at binding site position 1 to 8 (Table S2). We believe that disease-associated SNPs might be more prevalent in functional binding locations.

***iFeatures.*** Finally, iLearn is a newly presented toolkit that extracts a set of features, named as iFeatures, to describe DNA, RNA, and protein sequences [47, 48]. iLearn and iFeatures have been previously used in a variety of successful machine learning models in bioinformatics workflows [49, 50]. In this work, we applied iLearn to encode miRNA and mRNA target sequences in three different ways: (i) using a composition of *k*-spaced nucleotide pairs (Kmer), (ii) Dinucleotide-based Auto Covariance (DAC), and (iii) Pseudo Dinucleotide composition (PseDNC). This resulted in a set of 92 features (Table S2).

#### Statistical analysis

The difference between miRNA-target interactions, referred to as positive samples, and miRNA-non-target interactions, referred to as negative samples, was statistically investigated to acquire a better knowledge of target site features. Mann-Whitney *U* test was employed to determine the difference between the two population means in numerical characteristics (Table S8). For categorical variables, Chi-square ($\chi^2$) test was used to determine the association between two variables (Table S8).

#### Validation and Performance Metrics

***k*-fold Cross-validation (CV).** *k*-fold Cross-validation is a general approach for unbiased evaluation of a machine learning model's performance. This technique ensures that the model is evaluated on an independent dataset that is different from the training dataset. The method starts with a division of a dataset into *k* subsets called fold. For each fold, the model is trained with *k* - 1 subsets and validated with the remainder. The performance metric reported by *k*-fold cross-validation is the average of the results in each fold.

**Area Under the receiver operating characteristic Curve (AUC).** AUC is one of the most common evaluation metrics to evaluate a machine learning model for classification problems at various threshold settings. It is an area under the ROC curve plotted with True Positive Rate (TPR) and False Positive Rate (FPR), indicating the capability of the model to distinguish between classes. AUC values vary between 0 and 1, in which an excellent model achieves AUC near to 1. A model with AUC of 0.5 has no capability to separate between classes.

**Balanced accuracy (bACC).** bACC is another metric to evaluate the classification model. It is calculated as an average between True Positive Rate (TPR) or Sensitivity and True Negative Rate (TNR) or Specificity.

**Matthews Correlation Coefficient (MCC).** MCC is a statistical measure used for the assessment of binary classification models, evaluating the correlation between predicted and actual values. MCC values range from -1 to 1, with a perfect model receiving a score of 1. MCC is computed using the following formula:

$$MCC = \frac{TP \times TN - FP \times FN}{\sqrt{(TP + FP)(TP + FN)(TN + FP)(TN + FN)}}$$

Where TP, TN, FP, and FN refer to True positive, True negative, False positive, and False negative.

**Precision.** Precision is a commonly used metric for quantifying the performance of a model. It refers to the proportion of true positive sample predictions that were accurately predicted by the model.

**Recall, Sensitivity, or True Positive Rate (TPR).** Recall or Sensitivity refers to model capability to correctly detect positive samples. It is calculated as a total number of true positives divided by a total number of predicted positives.

**Specificity or True Negative Rate (TNR).** Specificity indicates a model capability to correctly predict negative samples as negative samples. It is calculated from a total number of true negatives divided by a total number of predicted negatives.

**F1-score.** F1-score. Using the harmonic mean, the F1-score combines the precision and recall of a classifier into a single statistic. Primarily, it is used to evaluate the performance of binary classification. High F1-score is only achievable if the classifier achieves both good precision and good recall.

### SUPPLEMENTARY RESULTS

#### Statistical analysis

In this work, Mann-Whitney *U* test and Chi-square test were performed to characterise the physical direct binding between miRNAs and target sites in mRNAs. The results of statistical analysis are summarised in Table S8. As hypothesised, the majority of features exhibit substantial differences in their means or proportions, and the majority are congruent with biological explanations. This consistency is associated with credence on the dataset's experimentally validated miRNA-target site interactions.

The outcome of the Chi-square test also confirmed that the percentage of 6-mer (the least effective site type) is greater in non-target samples, while the most effective type (8-mer) is the most prevalent in experimentally validated target samples. In samples where there are interactions between miRNA and target mRNAs (i.e., positive samples), the presence of nucleobase A at the initial position in mRNA and 3'-supplementary pairing is statistically more frequent than when the non-interactions are observed. Surprisingly, the ratios of SNPs and disease-associated SNPs in seed complementary sequences in mRNA do not vary significantly. Nevertheless, they are included in the feature selection process since the complicated pattern behind them might still improve the model performance.

Mann-Whitney *U* test demonstrates that miRNA-binding site interactions in positive samples are more energetically favourable than when non-interactions are found. Additionally, owing to complementary base pairs, 3'-supplementary bindings are more likely to occur in positive samples. In terms of site accessibility, the statistics indicate that a high level of accessibility can be implemented in the model due to the statistically higher AU content in binding sites and adjacent regions, the likelihood of binding sites being located near the start and the end of the 3'-UTR, and the lower energy required to make the site accessible. Another helpful indicator of binding site interaction is conservation. Given the functional significance of miRNA targeting sites, they are frequently conserved across species [10], which is consistent with our findings. 92 iFeature features [47, 48] were used to encode the miRNA and binding site nucleotide sequences in terms of Kmer, DAC, and PseDNC in this study. They demonstrate a pattern that is statistically distinct among target and non-target samples when all features are taken into account. As a result, we decided to include all iLearn features into a forward stepwise feature selection.

#### PRIMITI-TS Feature selection

In this study, a forward stepwise greedy feature selection [51-53] was employed to select the minimal, yet most effective combination of features. As a consequence, 22 out of 154 features were chosen to train a highly accurate model (Table S4). Two site type features, one binding stability features, one 3'-supplementary binding features, three site accessibility features, four conservation, two human genetic variation, and nine iFeature features were included in PRIMITI-TS's set of features. These features are used with XGBoost to train and validate the PRIMITI-TS model.

#### Performance of PRIMITI-TS

This work gathers the interactions between miRNAs and their target sites from various experimental data sources (Table S1). They are mixed and divided into training and test sets. However, the reported performance is often greater than the performance when applied to unseen experimental data. In this paper, we performed an experiment-split test in which the experimental sources for the training set are distinct from those for the test set in order to replicate data from a new experiment (Table S6). Although the performance varies amongst experiments, they all experience a considerable decrease in performance. Nonetheless, the model displays a significant level of generalisation across several datasets, with all AUC, bACC, F1, and MCC greater than 0.850, 0.800, 0.800, and 0.600 (Table S6). Consistency in Precision, Recall, and Specificity across experiments also suggest the robustness of the PRIMITI-TS model. Interestingly, even when training on human cells (Helwak *et al*. [17] and Kozar *et al*. [18]) and validating on vastly divergent *C. elegans* cells (Grosswendt *et al*. [16]), the model can still meet a satisfying performance with MCC of 0.628.

Around 300 distinct miRNAs are utilised to train the PRIMITI-TS model in this study. However, it only accounts for a small proportion of total human miRNAs as research suggests that there may be more than 2300 human miRNAs [54]. This raises a concern about the model's capability to operate with unseen miRNAs. A similar concern also applies to a number of transcripts in this study (3,331) to all protein-coding transcripts (~80,000) [55]. To validate the model performance with unseen miRNAs and/or mRNA, by randomly selecting 30% of miRNAs and mRNAs, we created (i) a dataset of unseen miRNA (ii) a dataset of unseen mRNA, and (iii) a dataset of unseen miRNA and mRNA. All datasets only show a small insignificant decrease in performance with AUC, bACC, F1, MCC, Precision, Recall, and Specificity values more than 0.900, 0.800, 0.800, 0.690, 0.880, 0.800, and 0.880 indicating that the model can still generalise well with unseen miRNA and mRNA (Table S7).

In this work, we introduce two new characteristics, iFeatures and SNPs, to further characterise miRNA-target site interactions. Previous miRNA-target prediction algorithms only encode miRNA and mRNA sequences using a basic technique, such as 1G (Guanine at the first nucleotide) and 5A (Adenine at the fifth nucleotide). PRIMITI-TS takes the lead by applying a sophisticated scheme accomplished by iLearn for encoding nucleic acid sequences into numerical features, called iFeatures. The other area that may be addressed is human genetic variations. We believe that the success of PRIMITI-TS is attributed to the addition of these features. The findings of a blind test and 5-fold, 10-fold, and 20-fold cross-validations (Table S10) indicate that iFeature features contribute considerably to miRNA-target interactions, with an MCC improvement of 0.708 to 0.788 in 5-fold cross-validation. Regrettably, there is no performance difference between models trained with and without SNPs features. It suggests that SNPs characteristics are unnecessary and may be omitted from the model.

Another major contribution of this work is a negative sample selection strategy. It was implemented to filter out low-quality negative samples, leading to a more reliable training dataset. To assess the performance improvement from a negative sample selection approach, models trained with a negative sample screened dataset and a dataset without negative sample selection were assessed under different cross-validation approaches and a blind test. The results exhibit significant improvements, with an increase in MCC from 0.773 to 0.796 in 5-fold cross-validation (Table S9).

### SUPPLEMENTARY DISCUSSION

Currently, a broad range of medicinal applications based on miRNA molecules have been created, many of them have undergone a clinical trial [56, 57]. That has attracted a huge interest in the domains of biomarkers, medicines, and pharmacological targets. However, a lack of comprehensive knowledge of miRNA activities hampered the development of therapeutic applications. Through direct binding with target mRNAs, miRNAs exert significant regulatory activity in molecular pathways, resulting in a post-transcriptional repression [30, 58, 59]. Thus, a variety of experimental approaches have been presented over the last decade, leading to a wide collection of confirmed miRNA-target mRNA interactions [8, 15-18, 22]. Nevertheless, the understanding of miRNA functions is far from completed, since the human genome contains more than 2,000 miRNAs and 20,000 protein-coding transcripts [54]. Hence, predictive-based computational strategies were created to bridge a knowledge gap and advance an understanding of miRNA functions. These tools may pave the way for the development of precise biomarkers and miRNA-based therapeutics that are more effective and have fewer side-effects.

In this study, to advance the definition of functional miRNA-target interactions, we have introduced three major improvements in terms of a negative sample selection to improve the quality of negative samples, coupled with the introduction of human genetic variation and iFeature characteristics. Relying on these improvements, PRIMITI-TS and PRIMITI-TM were introduced. First, the PRIMITI-TS model was developed and trained using a set of experimentally validated miRNA-target site interactions and non-interactions. Unlike other models, PRIMITI-TS was incorporated with a negative sample selection approach, which screened non-interactions using confirmed miRNA-target mRNA interactions. This procedure generates high-quality negative samples, which results in greater performance. Following that, 154 features were created to characterise miRNA-target site interactions and a minimal, but effective combination of 22 features was selected.

The performance of PRIMITI-TS was evaluated using an independent blind test and several cross-validation approaches. The performance levels and consistency across many validation situations demonstrate the robustness of in predicting potential miRNA-target site interactions. A further assessment on an external dataset also illustrates an excellent performance with an almost perfect score in recall, supporting the practicability of PRIMITI-TS for initial pre-screening of miRNA-target site interactions.

In this work, we had also introduced two sets of novel features, SNP features and iFeatures [47, 48]. An analysis was conducted to assess the effect of including the new features on the predictor's outcome, showing that iFeatures significantly contribute to PRIMITI-TS' predictive performance. This demonstrates that the enhanced, complex miRNA and mRNA sequence representation provided by iFeatures enables the description of RNA sequences from a variety of perspectives (e.g., DAC and PseDNC), which is beneficial for performance improvement.

Several real-world applications put a greater emphasis on the regulation of miRNA on target mRNA compared to the direct binding between miRNA and each target site [60-62]. Considering that a single mRNA may include several target sites, computational models provided a variety of approaches for combining the probability of miRNA-target sites in order to estimate the likelihood of miRNA-target mRNA repression activity. In this study, we used PRIMITI-TS to calculate the likelihood of miRNA-target site binding in each site. The landscape of miRNA-target site binding is represented by four features, the total of all probability values for all target sites, as well as the probabilities of the first, second, and third highest probability sites. PRIMITI-TM was created by XGBoost utilising these features to estimate the possibility of miRNA-induced post-transcriptional regulation on mRNA.

To provide an unbiased evaluation, an independent blind test and cross-validation procedures were performed. The significant AUROC, bACC, F1, and MCC values show that PRIMITI-TM performs well in terms of prioritising miRNA-target mRNA suppression. To demonstrate the practicability of PRIMITI-TM, we compared its performance to four state-of-the-art approaches, using experimentally confirmed miRNA-target mRNA interactions from Linsley's microarray [22] (Table 3). The results show that miRTarget outperforms other models, including PRIMITI-TM, in terms of overall performance, albeit at the expense of recall. In contrast, PRIMITI-TM can achieve the highest recall with a good precision, indicating the usability of the model for initial pre-screening. The similar pattern is evident in another validation of verified miRNA-target mRNA interactions gathered from the miRTarbase and Tarbase databases (Table 4). PRIMITI-TM exhibits the highest recall, which is about three times higher than miRTarget. Additionally, the PRIMITI-TM model also outperforms other competitors in terms of F1 and bACC overall performance. To enhance a prediction of potential miRNA-target mRNA interactions, we strongly suggest implementing PRIMITI-TM for initial pre-screening of miRNA-target mRNA interactions followed by the results of additional algorithms to increase the prediction's credibility.

In future works, the model prioritising miRNA-target interactions might be improved in different ways. The inability of existing models to determine the duplex structure of miRNA-target sites is one major limitation of current models. Up-to-date computational methods calculate the predicted miRNA-mRNA duplex by considering the most energetically and conformationally favourable RNA duplex structure. They include IntaRNA [63], RNAfold [64], RNAplfold [64], and dynamic programming used in PRIMITI and other state-of-the-art models. The prominent disadvantage is that the methods overlook the effect of other proteins on targeting mechanisms. For example, the binding at the ninth position of miRNA is considered unfavourable due to the structure of the silencing complex, a multiprotein complex. PRIMITI-TS aims to account for the effect of the protein complex by setting the penalty on binding. However, better structural prediction between miRNA and target site -- that takes the effect of protein complexes into consideration -- is needed to overcome this issue. Another promising area for further improvement is the representation of miRNA-target mRNA repression that is capable of identifying the effect of neighbouring sites. A number of studies has observed the cooperative activity between target sites, in which neighbouring sites may cooperatively repress mRNA expression under specific circumstances [65, 66]. It may be described by the ability of TNRC6, an important protein in miRNA-regulated repressional mechanisms playing a role in improving the stability of miRNA-target site interaction, to simultaneously bind with up to three complexes [65, 67]. In contrast to cooperative action, occlusive action can also be observed in some cases where the binding interrupts each other [35]. An insightful understanding of neighbouring site effects as well as better experimental techniques that can capture these effects are required in order to train a model that can facilitate cooperation between target sites.

**FIGURES**

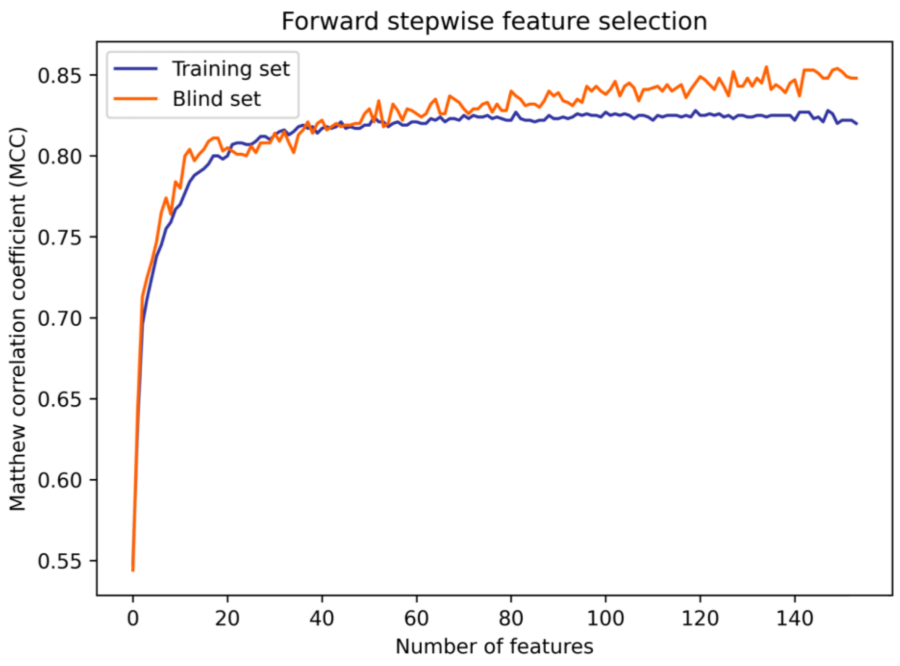

**Figure S1. Forward stepwise feature selection was employed to select the simplest but most effective combination of features.** Forward stepwise feature selection was implemented to minimise the dimensionality by selecting the optimal combination of features while avoiding overfitting. The approach begins with a set of zero features, evaluates each feature using a machine learning model, and then progressively adds the most beneficial feature in terms of Matthews correlation coefficient at each iteration. Each stage evaluated the model's performance using 10-fold cross-validation with an extreme gradient boosting classifier. The blind test line was shown only to analyse the behaviour of greedy feature selection, not being used to make a decision on the features themselves.

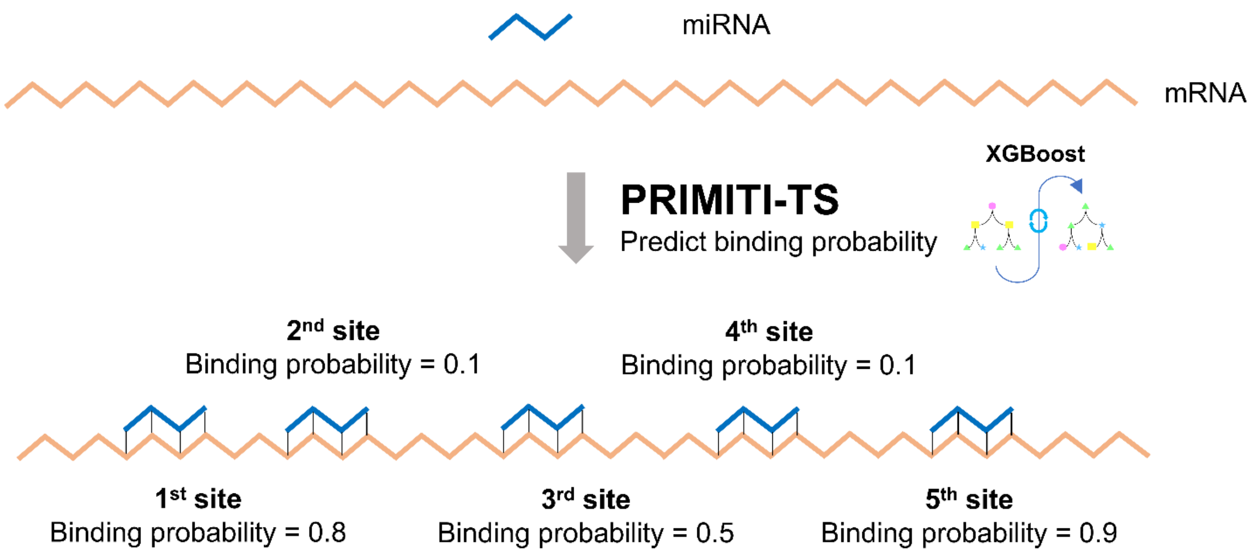

**4 Features**

| Summation | 1st Maximum probability | 2nd Maximum probability | 3rd Maximum probability |
|---|---|---|---|
| 2.4 | 0.9 | 0.8 | 0.5 |

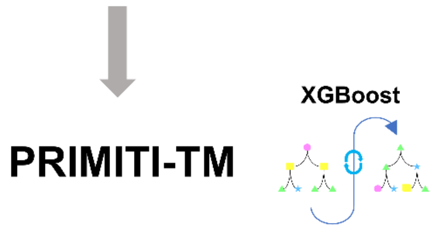

**Figure S2. PRIMITI-TM model is trained using four features derived from miRNA-target site functional binding probabilities estimated by PRIMITI-TS.** Initially, PRIMITI-TS was used to predict the functional binding between miRNA and canonical target site for each miRNA-mRNA interaction. All potential target sites with given probabilities will be transformed into four features: (i) The sum of all probabilities in all potential target sites, (ii) the probability in the target site with the highest probability, (iii) the probability in the target site with the second highest probability, (iv) the probability in the target site with the third highest probability. PRIMITI-TM model was trained using these four features.

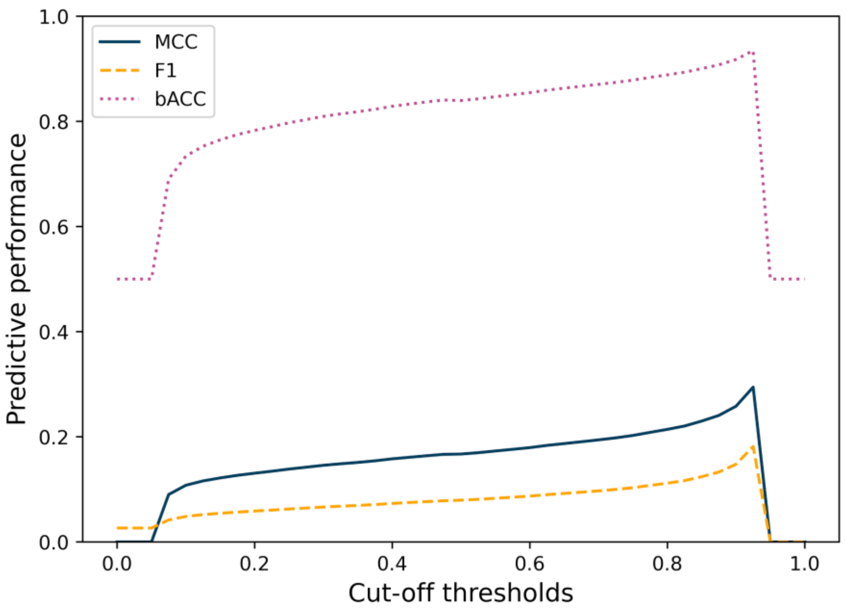

**Figure S3.** Performances of PRIMITI-TS on potential miRNA-target site interactions retrieved from Boudreau *et al.*'s HITS-CLIP under different cut-offs.

#### TABLES

**Table S1.** Analysed datasets in PRIMITI.

| Experimental type | Application | Reference | Data | |
|---|---|---|---|---|
| CLIP-seq | PRIMITI-TS training | (Helwak *et al.*, 2013) | 950 miRNA-target site interactions<br>117,512 miRNA-target site non-interactions | 6,190 miRNA-target site interactions<br>504,159 miRNA-target site non-interactions |
| CLIP-seq | PRIMITI-TS training | (Grosswendt *et al.*, 2014) | 883 miRNA-target site interactions<br>53,723 miRNA-target site non-interactions | |
| CLIP-seq | PRIMITI-TS training | (Kozar *et al.*, 2021) | 4,532 miRNA-target site interactions<br>373,224 miRNA-target site non-interactions | |
| CLIP-seq | PRIMITI-TS validation | (Boudreau *et al.*, 2014) | 98 miRNA-target site interactions<br>7,240 miRNA-target site non-interactions | |
| RNA-seq | PRIMITI-TM training | (Liu and Wang, 2019) | 2,351 miRNA-mRNA repression<br>6,749 miRNA-mRNA non-repression | |
| Microarray | PRIMITI-TM validation | (Linsley *et al.*, 2007) | 596 miRNA-mRNA repression<br>10,233 miRNA-mRNA non-repression | |

**Table S2.** 154 Features used to model miRNA-target site interactions in PRIMITI-TS. 22 Features were selected by a forward stepwise feature selection (highlighted in Bold).

| **Category** | **Description** | **Features** | **Sources / Tools** |
|---|---|---|---|
| Site types | A distinct subtype of the canonical target site is determined by the number of complementary bases and the presence of an adenine nucleobase at the initial position of the target site. In the case of a 6-mer, there exists an exact match of 6 nucleotides to the miRNA seed region (nt 2-7), whereas the 7-mer-m8 extends to the 8th nucleotide. The 7-mer-A1 shares similarities with the 6-mer but is distinguished by the inclusion of adenine ('A') at the first nucleotide of mRNA. On the other hand, the 8-mer encompasses both a perfect match at nt 2-8 and the presence of 'A' at the first nucleotide. | **6-mer** | Python |
| | | 7-mer-m8 | |
| | | 7-mer-A1 | |
| | | **8-mer** | |
| Binding stability | The stability of miRNA-target site duplex, represented by calculated energy from IntaRNA, an RNA-RNA interaction prediction program. | **Overall interaction energy** | IntaRNA (1), Python |
| | | Energy of hybridization | |
| | | Overall interaction energy of seed | |
| | | A at position 1 in mRNA | |
| 3'-supplementary binding | A supplementary binding occurred outside the seed region, preferably in supplementary region (nt 13-16), significantly contributing to the binding stability. | 3'-supplementary pairing in nucleotide position 13-16 | Python |
| | | Number of possible 3'-supplementary pairing | |
| | | **Number of possible complementary bases (positions 10 - 12)** | |
| | | Number of possible complementary bases (positions 13 - 16) | |
| | | Number of possible complementary bases (positions 17 +) | |
| | | Number of GC at miRNA positions 13-16 | |
| | | GG/CG/GC dinucleotide at miRNA position 13-16 | |
| Site accessibility | An AU content properties of target site and mRNA sequence. | Local 3'-UTR AU content (on binding site) | Python |
| | | Flank 3'-UTR AU content (± 30 nts) | |
| | | Flank 3'-UTR AU content (± 50 nts) | |

| | | | |
|---|---|---|---|
| | | **Global 3'-UTR AU content** | |
| | | Local 3'-UTR AU content (on binding site) / Global | |
| | | Flank 3'-UTR AU content (± 30 nts) / Global | |
| | | Flank 3'-UTR AU content (± 50 nts) / Global | |
| | A Location of target site in 3'-UTR of mRNA sequence. | Distance from start of 3'-UTR | Python |
| | | **Distance from terminal of 3'-UTR** | |
| | | Distance from start of 3'-UTR / UTR length | |
| | | **Distance from terminal of 3'-UTR / UTR length** | |
| | A degree of mRNA intramolecular interaction, which obstructs miRNA binding. | Energy required to make site accessible | Python |
| | | Probability to be accessible for mRNA | |
| Conservation | 30-way PhyloP.<br><br>A metric utilised to quantify a degree of evolutionary conservation of target site in multiple alignments of 30 vertebrate species. | 30-way PhyloP at position 1 | UCSC Genome Browser (2) |
| | | **30-way PhyloP at positions 2 - 7** | |
| | | 30-way PhyloP at position 8 | |
| | | 30-way PhyloP at positions 13 - 16 | |
| | 100-way PhyloP.<br><br>A metric utilised to quantify a degree of evolutionary conservation of target site in multiple alignments of 100 vertebrate species. | 100-way PhyloP at position 1 | UCSC Genome Browser (2) |
| | | 100-way PhyloP at positions 2 - 7 | |
| | | **100-way PhyloP at position 8** | |
| | | 100-way PhyloP at positions 13 - 16 | |
| | 30-way PhastCons.<br><br>A hidden Markov model-based method utilised to quantify a degree of evolutionary conservation of target site in multiple alignments of 30 vertebrate species. | 30-way PhastCons at position 1 | UCSC Genome Browser (2) |
| | | 30-way PhastCons at positions 2 - 7 | |
| | | **30-way PhastCons at position 8** | |

| | | | |
|---|---|---|---|
| | | 30-way PhastCons at positions 13 - 16 | |
| | 100-way PhastCons.<br><br>A hidden Markov model-based method utilised to quantify a degree of evolutionary conservation of target site in multiple alignments of 100 vertebrate species. | 100-way PhastCons at position 1 | UCSC Genome Browser (2) |
| | | **100-way PhastCons at positions 2 - 7** | |
| | | 100-way PhastCons at position 8 | |
| | | 100-way PhastCons at positions 13 - 16 | |
| Human genetic variation | A presence of single nucleotide polymorphisms (SNPs) on each location in a target site. | SNP at position 1 | miRNASNP-v3 (3) |
| | | SNP at position 2 | |
| | | SNP at position 3 | |
| | | SNP at position 4 | |
| | | SNP at position 5 | |
| | | SNP at position 6 | |
| | | SNP at position 7 | |
| | | SNP at position 8 | |
| | | SNP in seed | |
| | A presence of disease-related single nucleotide polymorphisms on each location in a target site. | Disease-related SNP at position 1 | miRNASNP-v3 (3) |
| | | Disease-related SNP at position 2 | |
| | | Disease-related SNP at position 3 | |
| | | Disease-related SNP at position 4 | |
| | | **Disease-related SNP at position 5** | |
| | | Disease-related SNP at position 6 | |
| | | Disease-related SNP at position 7 | |

| | | | |
|---|---|---|---|
| | | **Disease-related SNP at position 8** | |
| | | Disease-related SNP in seed | |
| iFeatures | A set of features for miRNA sequence representation using a composition of *k*-spaced nucleotide pairs (Kmer), dinucleotide-based auto covariance (DAC), pseudo dinucleotide composition (PseDNC). | **miRNA iFeature Kmer 1** | iFeature (4) |
| | | miRNA iFeature Kmer 2 | |
| | | miRNA iFeature Kmer 3 | |
| | | miRNA iFeature Kmer 4 | |
| | | miRNA iFeature Kmer 5 | |
| | | miRNA iFeature Kmer 6 | |
| | | miRNA iFeature Kmer 7 | |
| | | miRNA iFeature Kmer 8 | |
| | | miRNA iFeature Kmer 9 | |
| | | miRNA iFeature Kmer 10 | |
| | | miRNA iFeature Kmer 11 | |
| | | miRNA iFeature Kmer 12 | |
| | | miRNA iFeature Kmer 13 | |
| | | miRNA iFeature Kmer 14 | |
| | | miRNA iFeature Kmer 15 | |
| | | miRNA iFeature Kmer 16 | |
| | | miRNA iFeature DAC 1 | |
| | | miRNA iFeature DAC 2 | |
| | | miRNA iFeature DAC 3 | |

| | | |
|---|---|---|
| | | **miRNA iFeature DAC 4** |
| | | miRNA iFeature DAC 5 |
| | | miRNA iFeature DAC 6 |
| | | miRNA iFeature DAC 7 |
| | | miRNA iFeature DAC 8 |
| | | miRNA iFeature DAC 9 |
| | | miRNA iFeature DAC 10 |
| | | miRNA iFeature DAC 11 |
| | | miRNA iFeature DAC 12 |
| | | miRNA iFeature PseDNC 1 |
| | | miRNA iFeature PseDNC 2 |
| | | miRNA iFeature PseDNC 3 |
| | | miRNA iFeature PseDNC 4 |
| | | miRNA iFeature PseDNC 5 |
| | | miRNA iFeature PseDNC 6 |
| | | miRNA iFeature PseDNC 7 |
| | | miRNA iFeature PseDNC 8 |
| | | miRNA iFeature PseDNC 9 |
| | | **miRNA iFeature PseDNC 10** |
| | | **miRNA iFeature PseDNC 11** |
| | | miRNA iFeature PseDNC 12 |

| | | | |
|---|---|---|---|
| | | miRNA iFeature PseDNC 13 | |
| | | miRNA iFeature PseDNC 14 | |
| | | miRNA iFeature PseDNC 15 | |
| | | miRNA iFeature PseDNC 16 | |
| | | miRNA iFeature PseDNC 17 | |
| | | miRNA iFeature PseDNC 18 | |
| | A set of features for mRNA sequence representation using a composition of *k*-spaced nucleotide pairs (Kmer), dinucleotide-based auto covariance (DAC), pseudo dinucleotide composition (PseDNC). | mRNA iFeature Kmer 1 | iFeature (4) |
| | | mRNA iFeature Kmer 2 | |
| | | **mRNA iFeature Kmer 3** | |
| | | mRNA iFeature Kmer 4 | |
| | | mRNA iFeature Kmer 5 | |
| | | **mRNA iFeature Kmer 6** | |
| | | **mRNA iFeature Kmer 7** | |
| | | mRNA iFeature Kmer 8 | |
| | | mRNA iFeature Kmer 9 | |
| | | mRNA iFeature Kmer 10 | |
| | | mRNA iFeature Kmer 11 | |
| | | mRNA iFeature Kmer 12 | |
| | | mRNA iFeature Kmer 13 | |
| | | mRNA iFeature Kmer 14 | |
| | | mRNA iFeature Kmer 15 | |

| | | |
|---|---|---|
| | | mRNA iFeature Kmer 16 |
| | | mRNA iFeature DAC 1 |
| | | mRNA iFeature DAC 2 |
| | | mRNA iFeature DAC 3 |
| | | mRNA iFeature DAC 4 |
| | | mRNA iFeature DAC 5 |
| | | mRNA iFeature DAC 6 |
| | | mRNA iFeature DAC 7 |
| | | mRNA iFeature DAC 8 |
| | | mRNA iFeature DAC 9 |
| | | **mRNA iFeature DAC 10** |
| | | mRNA iFeature DAC 11 |
| | | mRNA iFeature DAC 12 |
| | | mRNA iFeature PseDNC 1 |
| | | mRNA iFeature PseDNC 2 |
| | | mRNA iFeature PseDNC 3 |
| | | mRNA iFeature PseDNC 4 |
| | | mRNA iFeature PseDNC 5 |
| | | mRNA iFeature PseDNC 6 |
| | | mRNA iFeature PseDNC 7 |
| | | mRNA iFeature PseDNC 8 |

| | | |
|---|---|---|
| | | mRNA iFeature PseDNC 9 |
| | | mRNA iFeature PseDNC 10 |
| | | mRNA iFeature PseDNC 11 |
| | | mRNA iFeature PseDNC 12 |
| | | mRNA iFeature PseDNC 13 |
| | | mRNA iFeature PseDNC 14 |
| | | mRNA iFeature PseDNC 15 |
| | | mRNA iFeature PseDNC 16 |
| | | mRNA iFeature PseDNC 17 |
| | | mRNA iFeature PseDNC 18 |

**Table S3.** The performance of PRIMITI-TS trained with 154 features under different classifiers under 10-fold cross-validation.

| **Algorithms** | **Area Under the Curve** | **Balanced Accuracy** | **F1** | **Matthews Correlation Coefficient** |
|---|---|---|---|---|
| **Extreme Gradient Boosting (XGBoost)** | **0.910 ± 0.001** | **0.910 ± 0.001** | **0.910 ± 0.001** | **0.821 ± 0.002** |
| Gradient Boosting | 0.906 ± 0.002 | 0.906 ± 0.002 | 0.906 ± 0.002 | 0.813 ± 0.004 |
| Support-Vector Classification (SVC) | 0.906 ± 0.001 | 0.906 ± 0.001 | 0.905 ± 0.001 | 0.812 ± 0.002 |
| Adaptive Boosting (ADABoost) | 0.896 ± 0.001 | 0.896 ± 0.001 | 0.896 ± 0.001 | 0.792 ± 0.002 |
| Random Forest (RF) | 0.895 ± 0.001 | 0.895 ± 0.001 | 0.894 ± 0.001 | 0.789 ± 0.002 |
| Neural Network | 0.894 ± 0.002 | 0.894 ± 0.002 | 0.894 ± 0.002 | 0.788 ± 0.004 |
| Extra Trees | 0.886 ± 0.001 | 0.886 ± 0.001 | 0.886 ± 0.001 | 0.770 ± 0.002 |
| J48 | 0.811 ± 0.003 | 0.811 ± 0.003 | 0.810 ± 0.003 | 0.621 ± 0.007 |

| | | | | |
|---|---|---|---|---|
| K-Nearest Neighbors (KNN) | 0.791 ± 0.001 | 0.791 ± 0.001 | 0.770 ± 0.002 | 0.593 ± 0.003 |
| Gaussian Processes | 0.773 ± 0.002 | 0.773 ± 0.002 | 0.752 ± 0.002 | 0.554 ± 0.004 |

**Table S4.** 22 Selected features used in PRIMITI-TS.

| Feature | Name | Feature | Name |
|---|---|---|---|
| 1 | 6-mer seed type | 12 | 8-mer seed type |
| 2 | miRNA iFeature DAC 4 | 13 | mRNA iFeature Kmer 4 |
| 3 | Overall interaction energy | 14 | Global 3’-UTR AU content |
| 4 | Distance from terminal of 3’-UTR | 15 | miRNA iFeature Kmer 1 |
| 5 | PhastCons 100-way nucleotide 2-7 | 16 | PhyloP 100-way nucleotide 8 |
| 6 | miRNA iFeature PseDNC 11 | 17 | PhastCons 30-way nucleotide 8 |
| 7 | Distance from terminal / 3’-UTR length | 18 | Disease-related SNPs nucleotide 5 |
| 8 | mRNA iFeature Kmer 3 | 19 | Disease-related SNPs nucleotide 8 |
| 9 | PhyloP 30-way nucleotide 2-7 | 20 | mRNA iFeature Kmer 7 |
| 10 | miRNA iFeature PseDNC 10 | 21 | mRNA iFeature DAC 10 |
| 11 | mRNA iFeature Kmer 6 | 22 | Number of possible complementary bases at nucleotide 10-12 |

**Table S5.** PRIMITI-TS validation on potential miRNA-target site interactions retrieved from Boudreau *et al.* HITS-CLIP in different cut-offs.

| Threshold | MCC | F1 | bACC | Precision | Recall | Specificity | TP | TN | FP | FN |
|---|---|---|---|---|---|---|---|---|---|---|
| **0.5** | 0.167 | 0.079 | 0.840 | 0.041 | 0.990 | 0.689 | 97 | 4990 | 2250 | 1 |
| **0.6** | 0.179 | 0.087 | 0.854 | 0.046 | 0.990 | 0.719 | 97 | 5207 | 2033 | 1 |
| **0.7** | 0.194 | 0.097 | 0.870 | 0.051 | 0.990 | 0.751 | 97 | 5436 | 1804 | 1 |
| **0.8** | 0.214 | 0.112 | 0.888 | 0.059 | 0.990 | 0.787 | 97 | 5697 | 1543 | 1 |
| **0.9** | 0.258 | 0.148 | 0.918 | 0.080 | 0.990 | 0.846 | 97 | 6122 | 1118 | 1 |

**Table S6.** Performance of PRIMITI-TS based on experiment-split validation on three experimental datasets.

| Training set | Validation set | AUC | bACC | F1 | MCC | Precision | Recall | Specificity |
|---|---|---|---|---|---|---|---|---|
| Helwak *et al.*<br>Grosswendt *et al.* | Ines *et al.* | 0.932 | 0.856 | 0.855 | 0.713 | 0.864 | 0.846 | 0.867 |
| Helwak *et al.*<br>Ines *et al.* | Grosswendt *et al.* | 0.895 | 0.813 | 0.803 | 0.628 | 0.845 | 0.766 | 0.859 |
| Grosswendt *et al.*<br>Ines *et al.* | Helwak *et al.* | 0.929 | 0.840 | 0.829 | 0.686 | 0.891 | 0.774 | 0.905 |

**Table S7.** The capability of PRIMITI-TS of predicting potential miRNA-target site interactions in unseen data.

| Datasets | AUC | bACC | F1 | MCC | Precision | Recall | Specificity |
|---|---|---|---|---|---|---|---|
| Unseen miRNA | 0.930 | 0.857 | 0.852 | 0.715 | 0.880 | 0.825 | 0.888 |
| Unseen mRNA | 0.957 | 0.891 | 0.889 | 0.782 | 0.907 | 0.872 | 0.910 |
| Unseen miRNA and mRNA | 0.924 | 0.844 | 0.838 | 0.691 | 0.872 | 0.807 | 0.881 |

**Table S8.** Statistical analysis of 154 Features. Categorical features were analysed with Chi-square test. Numerical features were analysed with Mann-Whitney U test.

| **Categorical features** | | | |
|---|---|---|---|
| **Features** | **Positive ratio** | **Negative ratio** | **P-value** |
| 6-mer site type | 0.094 | 0.560 | 0.0 |
| 7-mer-m8 site type | 0.088 | 0.177 | 1.196e-73 |
| 7-mer-A1 site type | 0.388 | 0.199 | 8.594e-298 |
| 8-mer site type | 0.430 | 0.064 | 0.0 |
| A at position 1 in mRNA | 0.518 | 0.241 | 0.0 |
| 3'-supplementary pairing in nucleotide position 13-16 | 0.701 | 0.576 | 1.727e-86 |
| GG/CG/GC dinucleotide at miRNA position 13-16 | 0.404 | 0.413 | 0.162 |
| SNP at position 1 in mRNA | 0.202 | 0.206 | 0.423 |
| SNP at position 2 in mRNA | 0.191 | 0.198 | 0.159 |
| SNP at position 3 in mRNA | 0.191 | 0.194 | 0.605 |
| SNP at position 4 in mRNA | 0.213 | 0.204 | 0.093 |
| SNP at position 5 in mRNA | 0.213 | 0.202 | 0.036 |
| SNP at position 6 in mRNA | 0.216 | 0.206 | 0.061 |
| SNP at position 7 in mRNA | 0.198 | 0.204 | 0.306 |
| SNP at position 8 in mRNA | 0.202 | 0.205 | 0.617 |
| SNP in seed complementary sequence in mRNA | 0.706 | 0.698 | 0.153 |
| Disease-related SNP at position 1 in mRNA | 0.014 | 0.012 | 0.227 |
| Disease-related SNP at position 2 in mRNA | 0.011 | 0.012 | 0.536 |
| Disease-related SNP at position 3 in mRNA | 0.009 | 0.011 | 0.203 |
| Disease-related SNP at position 4 in mRNA | 0.011 | 0.012 | 0.329 |
| Disease-related SNP at position 5 in mRNA | 0.014 | 0.012 | 0.118 |
| Disease-related SNP at position 6 in mRNA | 0.015 | 0.012 | 0.108 |
| Disease-related SNP at position 7 in mRNA | 0.013 | 0.012 | 0.909 |
| Disease-related SNP at position 8 in mRNA | 0.012 | 0.013 | 0.554 |
| Disease-related SNP in seed complementary sequence in mRNA | 0.070 | 0.068 | 0.637 |
| **Numerical features** | | | |
| **Features** | **Mean positive** | **Mean negative** | **P-value** |

| | | | |
|---|---|---|---|
| **Binding stability** | | | |
| Overall interaction energy | -13.238 | -9.150 | 0.0 |
| Energy of hybridization | -16.579 | -12.826 | 0.0 |
| Overall interaction energy of seed | -8.222 | -6.298 | 0.0 |
| **3’-supplementary binding** | | | |
| Number of possible 3’-supplementary pairing | 5.132 | 4.306 | 1.101e-76 |
| Number of possible complementary bases (positions 10 - 12) | 1.283 | 1.166 | 1.573e-16 |
| Number of possible complementary bases (positions 13 - 16) | 2.041 | 1.541 | 3.737e-142 |
| Number of possible complementary bases (positions 17 +) | 1.716 | 1.497 | 3.640e-25 |
| Number of GC at miRNA positions 13-16 | 1.889 | 2.047 | 2.093e-34 |
| **Site accessibility** | | | |
| Local 3’-UTR AU content (on binding site) | 0.549 | 0.540 | 1.741e-10 |
| Flank 3’-UTR AU content (± 30 nts) | 0.570 | 0.558 | 7.472e-17 |
| Flank 3’-UTR AU content (± 50 nts) | 0.569 | 0.559 | 2.415e-13 |
| Global 3’-UTR AU content | 0.568 | 0.567 | 0.001 |
| Local 3’-UTR AU content (on binding sit) / Global | 0.974 | 0.951 | 3.548e-13 |
| Flank 3’-UTR AU content (± 30 nts) / Global | 1.005 | 0.983 | 2.105e-22 |
| Flank 3’-UTR AU content (± 50 nts) / Global | 1.005 | 0.983 | 2.015e-22 |
| Distance from start of 3’-UTR | 1270.499 | 2210.626 | 0.0 |
| Distance from terminal of 3’-UTR | 1700.712 | 2242.989 | 1.612e-126 |
| Distance from start of 3’-UTR / UTR length | 0.449 | 0.493 | 8.040e-34 |
| Distance from terminal of 3’-UTR / UTR length | 0.537 | 0.499 | 4.564e-25 |
| Energy required to make site accessible | 3.341 | 3.676 | 2.059e-17 |
| Probability to be accessible for mRNA | 0.054 | 0.055 | 2.059e-17 |
| **Conservation** | | | |
| 100-way PhyloP at position 1 | 1.426 | 0.537 | 3.379e-288 |
| 30-way PhyloP at position 1 | 0.561 | 0.282 | 5.648e-237 |
| 100-way PhastCons at position 1 | 0.534 | 0.259 | 0.0 |

| | | | |
|---|---|---|---|
| 30-way PhastCons at position 1 | 0.584 | 0.317 | 0.0 |
| 100-way PhyloP at positions 2 - 7 | 1.825 | 0.582 | 0.0 |
| 30-way PhyloP at positions 2 - 7 | 0.677 | 0.352 | 0.0 |
| 100-way PhastCons at positions 2 - 7 | 0.583 | 0.265 | 0.0 |
| 30-way PhastCons at positions 2 - 7 | 0.604 | 0.319 | 0.0 |
| 100-way PhyloP at position 8 | 1.667 | 0.553 | 0.0 |
| 30-way PhyloP at position 8 | 0.595 | 0.288 | 2.065e-252 |
| 100-way PhastCons at position 8 | 0.553 | 0.259 | 0.0 |
| 30-way PhastCons at position 8 | 0.590 | 0.316 | 0.0 |
| 100-way PhyloP at positions 13 - 16 | 1.120 | 0.498 | 1.050e-268 |
| 30-way PhyloP at positions 13 - 16 | 0.516 | 0.318 | 4.287e-183 |
| 100-way PhastCons at positions 13 - 16 | 0.462 | 0.246 | 4.418e-277 |
| 30-way PhastCons at positions 13 - 16 | 0.529 | 0.308 | 0.0 |
| **iFeature features** | | | |
| miRNA iFeature Kmer 1 | 0.054 | 0.051 | 2.322e-06 |
| miRNA iFeature Kmer 2 | 0.040 | 0.047 | 7.300e-28 |
| miRNA iFeature Kmer 3 | 0.104 | 0.079 | 1.155e-204 |
| miRNA iFeature Kmer 4 | 0.050 | 0.049 | 1.450e-05 |
| miRNA iFeature Kmer 5 | 0.075 | 0.073 | 0.350 |
| miRNA iFeature Kmer 6 | 0.042 | 0.062 | 9.269e-145 |
| miRNA iFeature Kmer 7 | 0.020 | 0.019 | 0.008 |
| miRNA iFeature Kmer 8 | 0.057 | 0.082 | 0.0 |
| miRNA iFeature Kmer 9 | 0.049 | 0.050 | 0.006 |
| miRNA iFeature Kmer 10 | 0.070 | 0.067 | 0.0 |
| miRNA iFeature Kmer 11 | 0.068 | 0.075 | 0.002 |
| miRNA iFeature Kmer 12 | 0.094 | 0.069 | 8.633 |
| miRNA iFeature Kmer 13 | 0.071 | 0.049 | 2.833e-148 |
| miRNA iFeature Kmer 14 | 0.040 | 0.058 | 2.871e-174 |
| miRNA iFeature Kmer 15 | 0.100 | 0.098 | 0.485 |
| miRNA iFeature Kmer 16 | 0.067 | 0.072 | 0.006 |
| miRNA iFeature DAC 1 | -0.001 | -0.001 | 0.053 |
| miRNA iFeature DAC 2 | -0.000 | -0.000 | 1.505e-05 |

| | | | |
|---|---|---|---|
| miRNA iFeature DAC 3 | -1.431 | -1.020 | 1.038e-186 |
| miRNA iFeature DAC 4 | 0.179 | -0.164 | 3.433e-181 |
| miRNA iFeature DAC 5 | -0.000 | -0.000 | 0.007 |
| miRNA iFeature DAC 6 | -0.002 | -0.001 | 4.433e-143 |
| miRNA iFeature DAC 7 | 0.003 | 0.004 | 2.200e-24 |
| miRNA iFeature DAC 8 | -0.001 | -0.002 | 1.487e-124 |
| miRNA iFeature DAC 9 | -0.019 | -0.015 | 1.512e-09 |
| miRNA iFeature DAC 10 | -0.026 | -0.016 | 9.430e-12 |
| miRNA iFeature DAC 11 | -0.614 | -0.544 | 1.787e-33 |
| miRNA iFeature DAC 12 | -0.177 | -0.184 | 0.850 |
| miRNA iFeature PseDNC 1 | 0.036 | 0.035 | 0.015 |
| miRNA iFeature PseDNC 2 | 0.026 | 0.032 | 2.133e-46 |
| miRNA iFeature PseDNC 3 | 0.069 | 0.053 | 2.338e-160 |
| miRNA iFeature PseDNC 4 | 0.034 | 0.033 | 0.982 |
| miRNA iFeature PseDNC 5 | 0.049 | 0.049 | 0.007 |
| miRNA iFeature PseDNC 6 | 0.029 | 0.043 | 9.400e-139 |
| miRNA iFeature PseDNC 7 | 0.012 | 0.012 | 0.060 |
| miRNA iFeature PseDNC 8 | 0.039 | 0.057 | 5.572e-307 |
| miRNA iFeature PseDNC 9 | 0.033 | 0.034 | 1.422e-07 |
| miRNA iFeature PseDNC 10 | 0.045 | 0.044 | 0.005 |
| miRNA iFeature PseDNC 11 | 0.044 | 0.050 | 2.427e-23 |
| miRNA iFeature PseDNC 12 | 0.061 | 0.046 | 1.681e-77 |
| miRNA iFeature PseDNC 13 | 0.046 | 0.033 | 5.885e-117 |
| miRNA iFeature PseDNC 14 | 0.027 | 0.040 | 4.704e-171 |
| miRNA iFeature PseDNC 15 | 0.067 | 0.066 | 9.45e-06 |
| miRNA iFeature PseDNC 16 | 0.044 | 0.049 | 3.099e-12 |
| miRNA iFeature PseDNC 17 | 0.190 | 0.175 | 1.336e-248 |
| miRNA iFeature PseDNC 18 | 0.146 | 0.178 | 0.001 |
| mRNA iFeature Kmer 1 | 0.064 | 0.072 | 1.309e-08 |
| mRNA iFeature Kmer 2 | 0.062 | 0.050 | 8.102e-61 |
| mRNA iFeature Kmer 3 | 0.051 | 0.069 | 3.548e-168 |
| mRNA iFeature Kmer 4 | 0.058 | 0.059 | 0.155 |
| mRNA iFeature Kmer 5 | 0.079 | 0.073 | 1.376e-12 |
| mRNA iFeature Kmer 6 | 0.071 | 0.069 | 0.011 |

| | | | |
|---|---|---|---|
| mRNA iFeature Kmer 7 | 0.011 | 0.011 | 0.565 |
| mRNA iFeature Kmer 8 | 0.102 | 0.083 | 1.347e-109 |
| mRNA iFeature Kmer 9 | 0.041 | 0.052 | 1.143e-82 |
| mRNA iFeature Kmer 10 | 0.066 | 0.060 | 3.664e-18 |
| mRNA iFeature Kmer 11 | 0.042 | 0.059 | 8.289e-103 |
| mRNA iFeature Kmer 12 | 0.049 | 0.053 | 3.408e-16 |
| mRNA iFeature Kmer 13 | 0.064 | 0.053 | 2.140e-58 |
| mRNA iFeature Kmer 14 | 0.061 | 0.057 | 8.061e-06 |
| mRNA iFeature Kmer 15 | 0.087 | 0.086 | 0.551 |
| mRNA iFeature Kmer 16 | 0.092 | 0.094 | 0.654 |
| mRNA iFeature DAC 1 | -0.001 | -0.001 | 0.558 |
| mRNA iFeature DAC 2 | -0.000 | -0.000 | 1.553e-05 |
| mRNA iFeature DAC 3 | -1.170 | -0.935 | 6.420e-109 |
| mRNA iFeature DAC 4 | 0.050 | -0.045 | 1.759e-21 |
| mRNA iFeature DAC 5 | 0.000 | 0.000 | 1.161e-18 |
| mRNA iFeature DAC 6 | -0.001 | -0.001 | 1.072e-26 |
| mRNA iFeature DAC 7 | 0.003 | 0.004 | 3.061e-08 |
| mRNA iFeature DAC 8 | -0.001 | -0.002 | 3.334e-39 |
| mRNA iFeature DAC 9 | 0.010 | 0.010 | 0.470 |
| mRNA iFeature DAC 10 | -0.017 | -0.027 | 5.200e-14 |
| mRNA iFeature DAC 11 | -0.527 | -0.449 | 5.912e-86 |
| mRNA iFeature DAC 12 | -0.112 | -0.151 | 7.930e-17 |
| mRNA iFeature PseDNC 1 | 0.045 | 0.052 | 4.948e-10 |
| mRNA iFeature PseDNC 2 | 0.043 | 0.035 | 1.541e-56 |
| mRNA iFeature PseDNC 3 | 0.035 | 0.048 | 8.485e-177 |
| mRNA iFeature PseDNC 4 | 0.041 | 0.043 | 0.029 |
| mRNA iFeature PseDNC 5 | 0.055 | 0.051 | 9.411e-11 |
| mRNA iFeature PseDNC 6 | 0.050 | 0.049 | 0.012 |
| mRNA iFeature PseDNC 7 | 0.007 | 0.007 | 0.459 |
| mRNA iFeature PseDNC 8 | 0.071 | 0.058 | 7.119e-100 |
| mRNA iFeature PseDNC 9 | 0.029 | 0.037 | 1.187e-85 |
| mRNA iFeature PseDNC 10 | 0.045 | 0.040 | 3.008e-22 |
| mRNA iFeature PseDNC 11 | 0.028 | 0.040 | 4.443e-11 |
| mRNA iFeature PseDNC 12 | 0.033 | 0.037 | 1.120e-21 |

| | | | |
|---|---|---|---|
| mRNA iFeature PseDNC 13 | 0.045 | 0.037 | 1.241e-50 |
| mRNA iFeature PseDNC 14 | 0.043 | 0.041 | 0.001 |
| mRNA iFeature PseDNC 15 | 0.060 | 0.060 | 0.329 |
| mRNA iFeature PseDNC 16 | 0.066 | 0.067 | 0.564 |
| mRNA iFeature PseDNC 17 | 0.170 | 0.162 | 1.656e-79 |
| mRNA iFeature PseDNC 18 | 0.134 | 0.136 | 1.074e-08 |

**Table S9.** A performance of the PRIMITI-TS model without a negative sample selection strategy under 5-fold, 10-fold, and 20-fold cross-validation, and a blind test.

| **Methods** | **AUC** | **bACC** | **F1** | **MCC** | **Precision** | **Recall** | **Specificity** |
|---|---|---|---|---|---|---|---|
| **5-fold cross-validation** | | | | | | | |
| With negative sample selection | 0.959 ± 0.003 | 0.898 ± 0.006 | 0.897 ± 0.006 | 0.796 ± 0.012 | 0.900 ± 0.006 | 0.895 ± 0.008 | 0.900 ± 0.006 |
| Without negative sample selection | 0.954 ± 0.004 | 0.886 ± 0.004 | 0.886 ± 0.004 | 0.773 ± 0.008 | 0.887 ± 0.003 | 0.885 ± 0.007 | 0.887 ± 0.004 |
| **10-fold cross-validation** | | | | | | | |
| With negative sample selection | 0.959 ± 0.005 | 0.899 ± 0.007 | 0.899 ± 0.006 | 0.798 ± 0.013 | 0.901 ± 0.013 | 0.896 ± 0.010 | 0.901 ± 0.015 |
| Without negative sample selection | 0.956 ± 0.007 | 0.888 ± 0.013 | 0.888 ± 0.013 | 0.775 ± 0.026 | 0.887 ± 0.014 | 0.888 ± 0.017 | 0.887 ± 0.015 |
| **20-fold cross-validation** | | | | | | | |
| With negative sample selection | 0.960 ± 0.009 | 0.903 ± 0.017 | 0.903 ± 0.018 | 0.807 ± 0.035 | 0.906 ± 0.018 | 0.901 ± 0.027 | 0.906 ± 0.020 |
| Without negative sample selection | 0.956 ± 0.012 | 0.891 ± 0.020 | 0.891 ± 0.019 | 0.782 ± 0.040 | 0.893 ± 0.026 | 0.889 ± 0.017 | 0.893 ± 0.027 |
| **Blind test** | | | | | | | |
| With negative sample selection | 0.966 | 0.904 | 0.903 | 0.809 | 0.915 | 0.892 | 0.917 |
| Without negative sample selection | 0.959 | 0.899 | 0.898 | 0.798 | 0.906 | 0.890 | 0.908 |

**Table S10.** Predictive performance of PRIMITI-TS model with and without iFeature and SNPs features.

| **Methods** | **AUC** | **bACC** | **F1** | **MCC** | **Precision** | **Recall** | **Specificity** |
|---|---|---|---|---|---|---|---|
| **5-fold cross-validation** | | | | | | | |
| With all features | 0.959 ± 0.003 | 0.898 ± 0.006 | 0.897 ± 0.006 | 0.796 ± 0.012 | 0.900 ± 0.006 | 0.895 ± 0.008 | 0.900 ± 0.006 |
| Without iFeature | 0.927 ± 0.010 | 0.853 ± 0.005 | 0.853 ± 0.005 | 0.707 ± 0.010 | 0.856 ± 0.005 | 0.849 ± 0.007 | 0.857 ± 0.005 |
| Without SNPs | 0.959 ± 0.002 | 0.897 ± 0.004 | 0.897 ± 0.004 | 0.794 ± 0.007 | 0.898 ± 0.003 | 0.896 ± 0.006 | 0.898 ± 0.002 |
| Without iFeature & SNPs | 0.926 ± 0.005 | 0.853 ± 0.003 | 0.852 ± 0.003 | 0.705 ± 0.006 | 0.855 ± 0.003 | 0.850 ± 0.006 | 0.855 ± 0.004 |
| **10-fold cross-validation** | | | | | | | |
| With all features | 0.959 ± 0.005 | 0.899 ± 0.007 | 0.899 ± 0.006 | 0.798 ± 0.013 | 0.901 ± 0.013 | 0.896 ± 0.010 | 0.901 ± 0.015 |
| Without iFeature | 0.928 ± 0.010 | 0.856 ± 0.010 | 0.855 ± 0.010 | 0.712 ± 0.020 | 0.857 ± 0.013 | 0.853 ± 0.014 | 0.858 ± 0.015 |
| Without SNPs | 0.959 ± 0.006 | 0.897 ± 0.007 | 0.897 ± 0.007 | 0.794 ± 0.014 | 0.899 ± 0.012 | 0.895 ± 0.010 | 0.899 ± 0.013 |
| Without iFeature & SNPs | 0.928 ± 0.009 | 0.855 ± 0.010 | 0.855 ± 0.010 | 0.711 ± 0.020 | 0.857 ± 0.012 | 0.853 ± 0.014 | 0.858 ± 0.013 |
| **20-fold cross-validation** | | | | | | | |
| With all features | 0.960 ± 0.009 | 0.903 ± 0.017 | 0.903 ± 0.018 | 0.807 ± 0.035 | 0.906 ± 0.018 | 0.901 ± 0.027 | 0.906 ± 0.020 |
| Without iFeature | 0.929 ± 0.012 | 0.856 ± 0.015 | 0.856 ± 0.016 | 0.713 ± 0.030 | 0.857 ± 0.019 | 0.855 ± 0.025 | 0.857 ± 0.023 |
| Without SNPs | 0.960 ± 0.010 | 0.902 ± 0.014 | 0.902 ± 0.015 | 0.805 ± 0.029 | 0.905 ± 0.017 | 0.899 ± 0.024 | 0.906 ± 0.019 |

| | | | | | | | |
|---|---|---|---|---|---|---|---|
| Without iFeature & SNPs | 0.929 ± 0.013 | 0.858 ± 0.016 | 0.858 ± 0.017 | 0.716 ± 0.032 | 0.860 ± 0.018 | 0.856 ± 0.026 | 0.860 ± 0.021 |
| **Blind test** | | | | | | | |
| With all features | 0.966 | 0.904 | 0.903 | 0.809 | 0.915 | 0.892 | 0.917 |
| Without iFeature | 0.933 | 0.863 | 0.862 | 0.726 | 0.870 | 0.853 | 0.872 |
| Without SNPs | 0.966 | 0.906 | 0.904 | 0.812 | 0.919 | 0.891 | 0.921 |
| Without iFeature & SNPs | 0.933 | 0.865 | 0.864 | 0.731 | 0.874 | 0.854 | 0.877 |
